# Recurrent mutations drive rapid HIV escape from two broadly neutralizing antibodies *in vivo*

**DOI:** 10.1101/2025.08.29.673185

**Authors:** Elena V. Romero, Abigail E. Clyde, Elena E. Giorgi, Dylan H. Westfall, Walker Azam, Megan L. Taylor, Marina Caskey, Alison F. Feder, Lillian B. Cohn

**Affiliations:** Department of Genome Sciences, University of Washington, Seattle, WA, USA; Vaccine and Infectious Diseases Division, Fred Hutchinson Cancer Center, Seattle, WA, USA; Laboratory of Molecular Immunology, The Rockefeller University, New York, NY, USA; Division of Public Health Sciences, Fred Hutch Cancer Center, Seattle, WA, USA; Department of Global Health, University of Washington, Seattle, WA, USA

## Abstract

Broadly neutralizing antibodies (bNAbs) show promise for HIV treatment and prevention, but are vulnerable to resistance evolution. Comprehensively understanding *in vivo* viral escape from individual bNAbs is necessary to design bNAb combinations that will provide durable responses. We characterize viral escape from two such bNAbs, 10-1074 and 3BNC117, using deep, longitudinal sequencing of full length HIV envelope (*env*) genes from study participants treated with bNAb monotherapy. Improved sequencing depth and computational evolutionary analyses permit us to identify *in vivo* routes and parallelism underlying HIV escape from each bNAb, providing new insights into this evolutionary process. We find that 10-1074 escape is restricted to a small number of previously documented pathways seen across participants, but these escape mutations 1) emerge via extensively recurrent mutation, 2) are not equally preferred, and 3) can pre-exist at low frequency in intra-host viral populations before therapy, although their detection does not predict rebound timing. In contrast, 3BNC117 escape follows background-specific patterns in which specific escape mutations present in one intra-host population rarely emerge or spread in other populations, except among highly related viruses. Despite this, 3BNC117 escape mutations often still emerge recurrently within their host. Our findings map longitudinal *in vivo* antibody escape across 20 diverse clade B HIV intra-host populations and reveal clinically relevant resistance dynamics that highlight how combination bNAb therapies will need to contend with extensively recurring escape mutations and dependence on genetic background.

**Significance Statement:** Using recently developed techniques that capture viral genetic diversity and associations between mutations at depth, we deeply sequenced HIV from two clinical trials of broadly neutralizing antibody (bNAb) monotherapies, 3BNC117 and 10-1074. We computationally characterized HIV populations longitudinally with unprecedented resolution as they escaped these therapies in people living with HIV. Intra-host tracking of individual HIV genetic backgrounds reveals extensively recurrent mutations driving escape and suggests that HIV escape routes from certain bNAbs can depend sensitively on the genetic background of the virus. Our findings highlight the difficulties in evaluating pre-treatment resistance, provide an analysis blueprint for future trials, and inform the design of emerging combination antibody therapies to maximize the likelihood of durable efficacy.

## Introduction

Combination antiretroviral therapy (ART) has revolutionized the treatment and prevention of HIV-1 infection. However, ART does not eradicate established infection (Chun et al., 1997; Cohn et al., 2020) and worldwide HIV-1 incidence rates remain high (UNAIDS Data 2024 | UNAIDS, 2024). Thus, the search for novel preventive and therapeutic interventions remains a global high priority. Broadly neutralizing antibodies (bNAbs) have emerged as a possible new therapeutic and preventive strategy because antibodies allow for infrequent dosing, directly target multiple viral epitopes, and have the potential to harness host immune responses via their Fc effector domains (Bournazos et al., 2017). However, their clinical development has been hindered in part because bNAbs are vulnerable to escape by HIV-1 variants.

*In vivo*, HIV rapidly escapes bNAb monotherapies through the generation of resistance (i.e. escape) mutations (Caskey et al., 2015, 2017; Lynch et al., 2015; Stephenson et al., 2021). However, combining bNAbs into co-administered cocktails has begun to yield promising results (Eron et al., 2024; Gaebler et al., 2022; Julg et al., 2024), mirroring the therapeutic development of ART cocktails in the 1990s. Different bNAbs target different sites of the HIV Env protein spike, including the CD4 binding site (CD4bs), the V1V2-glycan epitope, the V3-loop region, gp120-gp41 interface and the membrane-proximal external region (MPER). Multiple bNAbs applied simultaneously thus challenge the virus combinatorially, raising the genetic barrier to full resistance (Julg et al., 2024; Mahomed et al., 2022; Mendoza et al., 2018; Sneller et al., 2022; Mayer et al., 2022; Wagh et al., 2016). Recent work has shown that viral escape from pooled antibodies can be well described by investigating the action of each antibody component independently then employing an additive model of their effects (Dingens et al., 2019; Yu et al., 2022). Realizing combination bNAb therapy’s potential as a long-term treatment, prevention, and possibly even cure, of HIV will likely therefore rely on comprehensively characterizing viral escape from individual bNAbs to design optimal combinations (LaMont et al., 2022). However, two major challenges exist.

First, the diversity of viral escape routes from individual bNAbs is incompletely understood. *In vitro* approaches like deep mutational scanning (DMS) permit the systematic characterization of viral escape phenotypes against specific bNAbs by measuring the effects of all single mutations away from a given reference strain (Bricault et al., 2019; Dingens et al., 2019; LaBranche et al., 2018; Radford & Bloom, 2025a; Schommers et al., 2020).

However, it remains unclear if these represent the full range of escape trajectories achieved *in vivo* and how these results generalize to new viral genetic backgrounds. For example, Radford et al. recently found that two divergent reference envelope strains from different subtypes achieved 3BNC117 escape via different mutations, suggesting that escape from some bNAbs may be dependent on the genotypic context in which mutations arise (Radford & Bloom, 2025a). This may limit the utility of DMS for predicting escape variants clinically, as participant-specific viruses are genetically distinct from DMS references. Therefore, profiling escape mutations *ex vivo* from diverse HIV populations treated with bNAbs is necessary to determine escape pathways in practice, assess their dependence on genetic background, and uncover escape-associated genetic variation like insertions and deletions which are not generally evaluated in DMS experiments.

Second, while it is clear that most bNAb monotherapy-treated HIV populations escape treatment, it is unclear how *recurrently* these populations produce escape mutations *in vivo*. Understanding this degree of evolutionary parallelism can provide insight into the simultaneous challenges necessary to durably suppress HIV. For example, recognition of parallelism driving escape from ART monotherapy helped scientists establish that multiple distinct therapies needed to be combined to effectively limit resistance (Coffin, 1995). Similarly, profiling the degree of parallel evolution during bNAb escape could improve estimates of how many components would be needed for combination therapies to achieve durable suppression. Parallelism can provide this insight because it serves as a proxy for how easily viral populations produce escape mutations.

Parallel escape can emerge because many different mutations confer escape, genomes mutate at a high rate, and/or because the population size of those viruses that can produce the mutations is large, all of which are likely to be important here. Although previous bNAb monotherapy studies have observed that intra-host viral populations also appear to escape via multiple mutations concurrently (Caskey et al., 2015, 2017; Williamson et al., 2025), such parallelism has not been investigated comprehensively due to an incomplete understanding of accessible escape routes and because mutations that are produced recurrently within a population are not easily recognized as such.

We therefore sought to understand the *in vivo* routes and parallelism of HIV escape from two broadly neutralizing antibodies, 10-1074 and 3BNC117, which target the V3 glycan and CD4 binding site, respectively.

Previous attempts to address these questions have been constrained by limited sequencing depth using low-throughput single genome analysis methods (Caskey et al., 2015, 2017; Schoofs et al., 2016). While deep sequencing approaches can characterize *in vivo* viral evolution with high resolution, they rely on short reads where information about viral genetic background is lost (Caskey et al., 2017; Zanini et al., 2015). We therefore performed long-read deep sequencing on longitudinal plasma samples from two bNAb monotherapy clinical trials of 10-1074 and 3BNC117. This approach, called SMRT-UMI, permitted substantially deeper sequencing than previously employed approaches while also retaining critical information about the genetic backgrounds on which resistance mutations occur (Westfall et al., 2024). These advances allowed us to develop a computational pipeline incorporating novel strategies to screen for escape mutations and parallelism, revealing new insights into *in vivo* therapeutic escape for both bNAbs and serving as a blueprint for future investigations of viral intra-host evolution.

## Results

### High resolution sequencing yields fully resolved *env* haplotypes

To collect high resolution, linkage-resolved sequences from HIV populations escaping bNAbs, we performed long-read, high throughput, deep sequencing (SMRT-UMI) of full-length HIV envelope (Fig. 1a) (Westfall et al., 2024). Briefly, we attach a unique molecular identifier (UMI) to viral cDNA prior to amplification. We then perform PCR to amplify HIV envelope and use a long-read sequencing approach to capture the full-length HIV *env* gene and its linked UMI. This method is unique because it permits deeper sequencing than standard single genome approaches, which increases statistical power to discover putative resistance mutations, while still preserving linkage information via UMI-based cDNA tagging to control for *in vitro* recombination.

**Figure 1:**
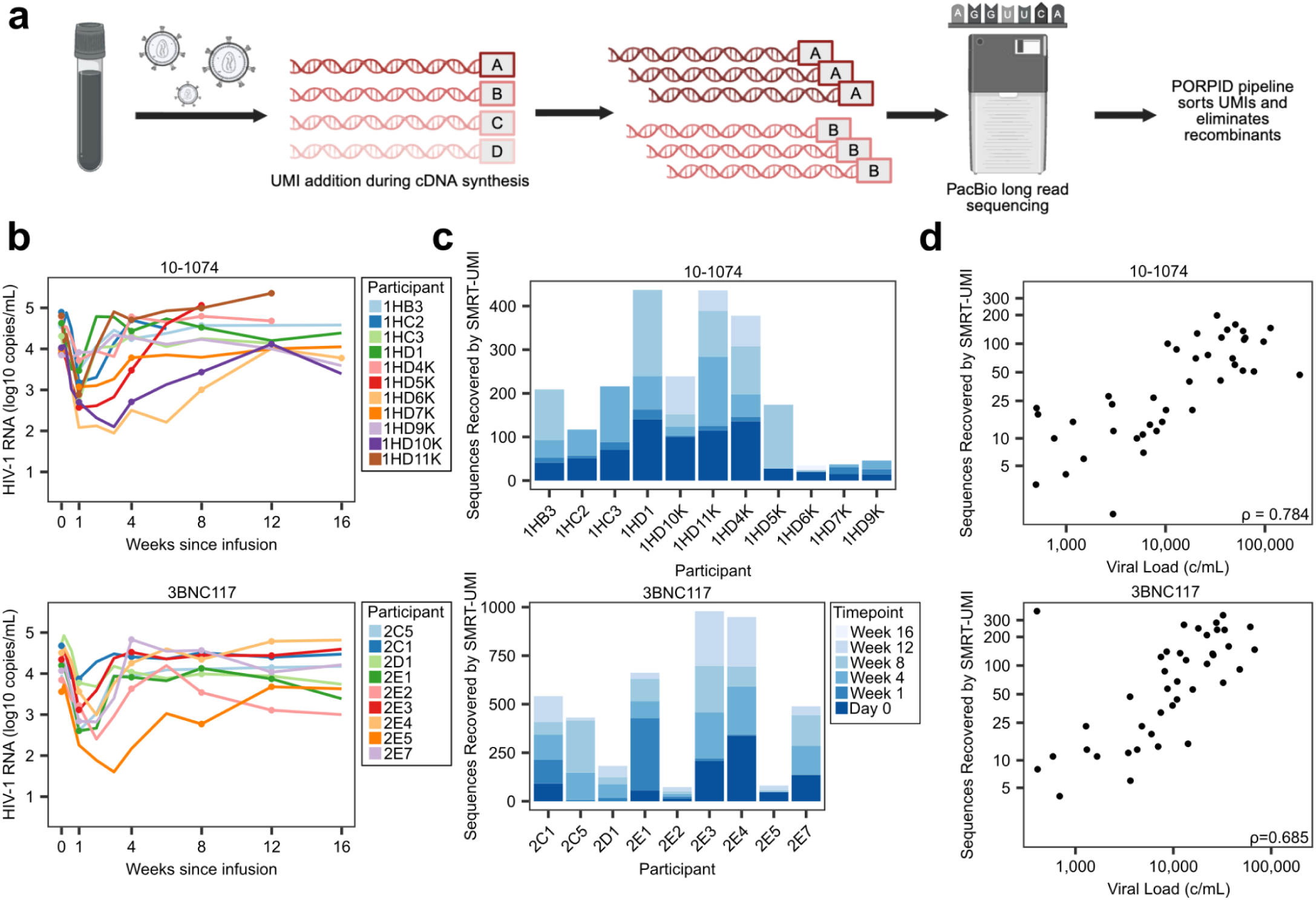
High-resolution sequence recovery of samples from human clinical trials testing two broadly neutralizing antibody monotherapies infused during chronic HIV infection. (**a**) Schematic representation of SMRT-UMI sequencing protocol. (**b**) Viral load graphs following 10-1074 (top) or 3BNC117 (bottom) infusion from NCT 02511990 and NCT 02018510, respectively. Each color represents a participant and dots indicate timepoints sequenced (see Supp. Tables 1,2 for complete sampling scheme) (**c**) Number of sequences recovered per participant using SMRT-UMI, separated by time point. Sequences recovered in 10-1074 treated samples (top) and 3BNC117 treated samples (bottom). (**d**) The number of sequences recovered using SMRT-UMI is positively correlated with viral load (copies/mL) in the 10-1074 treated samples (top) and 3BNC117 treated samples (bottom). Spearman ρ=0.784, 𝑝= 1.352 × 10 ^−6^ (top); ρ = 0. 685, 𝑝 = 7. 90 × 10 (bottom).

To understand viral escape from bNAbs targeting distinct epitopes, we applied this sequencing approach to samples from two clinical trials. 10-1074 was tested in NCT 02511990 and 3BNC117 was tested in NCT 02018510. Both studies involved a single administration of the respective antibody to viremic participants living with HIV (PLWH) (Fig. 1b) (Caskey et al., 2015, 2017). Both 10-1074 and 3BNC117 reduced plasma viremia by 1-3 logs in the majority of participants (Fig 1b). Viral loads rebounded to pre-treatment levels by weeks 4-12 in both trials, driven by a combination of declining antibody concentrations and the emergence of resistant variants. We sequenced samples from timepoints day 0, week 1, and week 4 after antibody administration, and at later timepoints in a subset of individuals (Fig. 1d, Supp. Tables 1,2 for full sampling scheme).

In total, we recovered 2324 *env* sequences from 11 study participants treated with 10-1074 (median 40; range: 1-198), and 4405 full length *env* sequences from 9 study participants treated with 3BNC117 (median 68; range: 4-373), all of which were Clade B (Fig. 1c). Importantly, this included sequencing of a transmission pair treated with 3BNC117 (2E4 and 2E5), which provided an opportunity to examine *in vivo* mutational escape pathways in two highly related intra-host populations. Two timepoints (2E5 week 1 and week 4, 17 sequences) were sequenced using conventional single genome sequencing methods due to low viral loads and limited previously reported data (Caskey et al., 2015). Relative to the original single genome sequencing studies, our sequencing depth increased 3.5x for 10-1074 (2324 vs 657 sequences) and 7.5x for 3BNC117 (4388 vs 573 sequences) (Caskey et al., 2015, 2017; Schoofs et al., 2016). This expanded dataset provided the statistical power and temporal resolution to quantify intrahost evolution, track competing viral lineages, and identify resistant variants present at low frequency that were not observable in earlier datasets.

Recovery of sequences varied by participant and correlated with viral load (Fig. 1d; 10-1074 Spearman ρ = 0. 784, 𝑝 = 1. 352 × 10^−6^; 3BNC117 ρ = 0. 685, 𝑝 = 7. 90 × 10^−7^). For downstream analyses, we aligned the filtered sequences by participant along with previously recovered SGA sequences and confirmed absence of donor-to-donor contamination (Supp. Fig. 1, 2, Supp. Tables 3,4, Caskey et al., 2015, 2017; Schoofs et al., 2016).

### Viruses escaping 10-1074 show both novel and previously reported resistance mutations

To assess the full range of escape pathways observed during 10-1074 treatment, we investigated HIV amino acid (AA) sequence changes. We identified variable loci in 10-1074 treated participants where the day 0 consensus AA decreased by at least 40% in frequency after bNAb administration (see Materials & Methods). In concordance with previous shallower single genome sequencing results (Caskey et al., 2017), we identified large and consistent AA changes at loci N332, S334 and D/N325 after 10-1074 infusion which partially reverted in some participants by the end of the trial, potentially indicating a fitness cost of escape (Fig. 2a-d, Supp. Fig. 3, Supp. Table 5). Deeper sequencing revealed additional AA identities in 7/9 participants, including 332T which was newly observed at 10% frequency in participant 1HB3 at week 4 (Fig. 2a), and 332H which has not previously been linked to *in vivo* escape. Mutations at N332 and S334 eliminate the N332 glycan bound by 10-1074 and appear to be independent of viral genetic background. The one exception was participant 1HD9K whose HIV population had a pre-existing escape mutation at another 10-1074 contact site, D/N325E, potentially lessening the selective benefit of N332 glycan loss.

**Figure 2.**
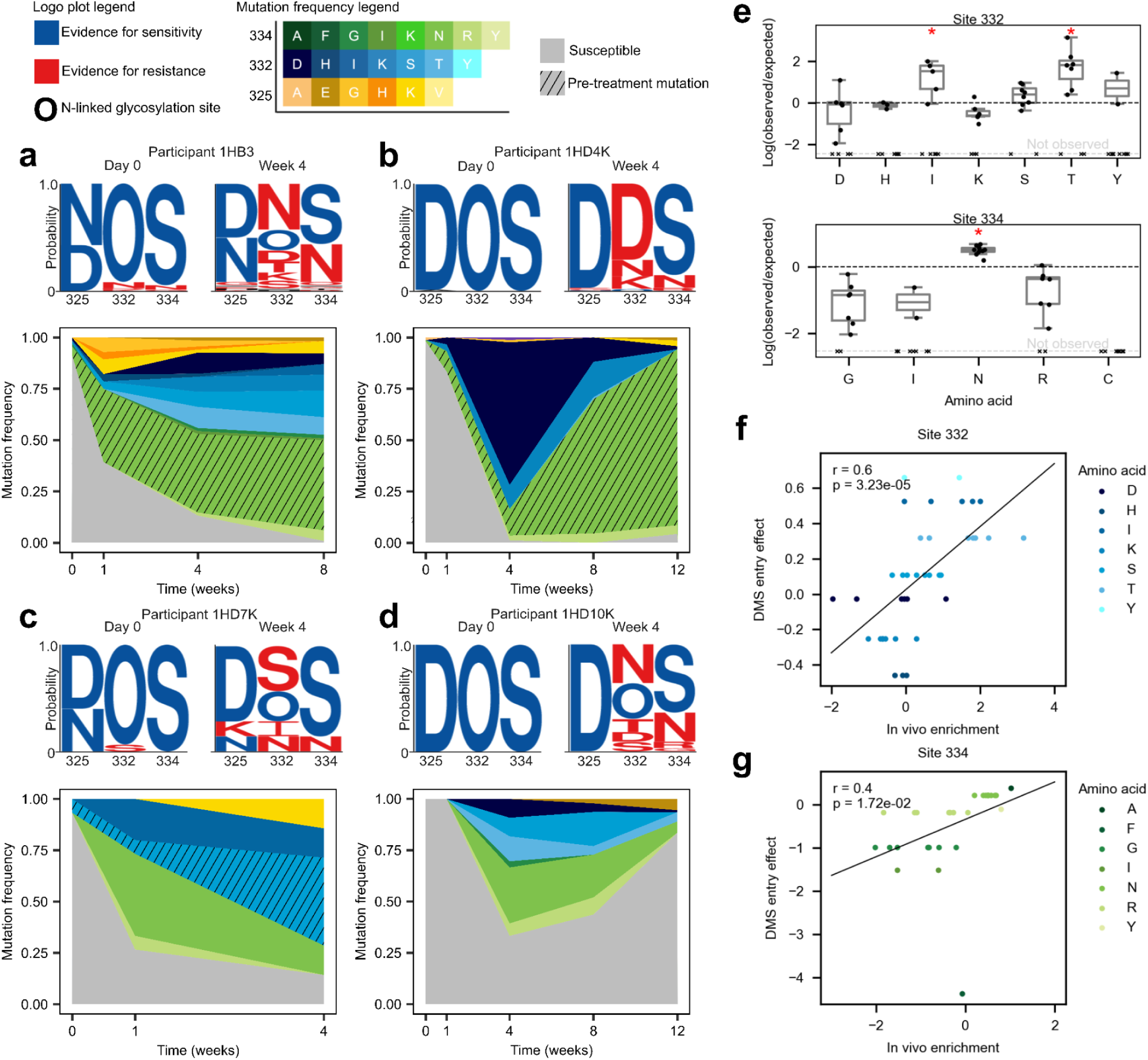
Dynamics and relative enrichments of viral escape mutations following 10-1074 treatment. **(a-d)** (top) Logo plots of amino acids (AAs) at loci 325, 332, and 334 are shown for pre-treatment time point (day 0) and the first post-nadir time point (week 4) in select 10-1074 treated participants. AAs in red or blue have previously been reported to confer resistance or sensitivity to 10-1074, respectively (Bricault et al., 2019; Caskey et al., 2017; Dingens et al., 2019; Radford & Bloom, 2025a). (bottom) Frequency plots of escape mutations over time are shown, with each mutation indicated by color and hatching indicating those escape mutations discovered pre-treatment. Additional participants are shown in supplemental figure 3. **(e)** Observed mutational frequencies in the trial data were compared to predicted mutational frequencies calculated based on the probability of the day 0 ancestral codon mutating to a given AA (See Materials & Methods) at loci 332 & 334. Ratios of log(Observed/Expected) > 0 indicate an enrichment above neutral expectation. Each data point represents the comparison for a single participant. Hatch marks on the “Not observed” line indicate participants in which the change was expected by mutation, but not observed. Significance was determined via binomial tests with a Bonferroni correction (significance level α = 3. 1 × 10^−3^, 𝑝 = 6. 6 × 10^−9^, 1. 2 × 10^−4^, 7. 5 × 10^−35^ for N332T, N332I, and S334N, respectively). **(f,g)** *In vivo* enrichment, log(Observed/Expected), for each AA sampled in each participant, plotted against cell entry effects for TZM-bl cells as determined by deep mutational scanning of the clade B TRO11 HIV Env protein (Radford & Bloom, 2025a) for sites 332 and 334. Pearson 𝑟 = 0.6, ^−5^ (site 332); 𝑟 = 0. 4, ^−2^ (site 334).

To look for additional putative escape mutations, we identified all loci that showed ≥ 40% day 0 consensus AA loss across 3 or more participants (Supp. Table 5). Notably, V4 locus 406 showed AA changes in 4 individuals, although the locus may not be comparable across all individuals due to alignment challenges in this hypervariable region (Supp. Table 5, Supp. Fig. 5). While, to our knowledge, this is the first report of locus 406 being associated with resistance to 10-1074 *in vivo*, DMS has linked this locus to escape from PGT121, another V3-targeting bNAb, *in vitro* previously (Dingens et al., 2019). We also applied the Marginal Path Likelihood (MPL), a method that can detect sites under selection from longitudinal samples without relying on parallelism across participants (Sohail et al., 2021). We found broadly concordant results, with sites 325, 332 and 334 detected as under selection independently in many different participants, although due to the conservative run parameters (see Methods), we did not detect selected sites in all participants (Supp. Table 6). MPL also detected that nearby site 328 was under positive selection in 1HD4K, a change that was not observed in any other participant.

We compared our results to those mutations identified *in vitro* by Radford and Bloom in their DMS of 10-1074 escape mutations in two divergent HIV backgrounds (Radford & Bloom, 2025a).There was a positive correlation between DMS escape score and the maximum frequency a mutation achieved *in vivo* (Supp. Fig. 4). The strongest hits (at N332, S334 and D/N325) appeared both *in vivo* and across both *in vitro* genetic backgrounds (Supp. Fig. 4). However, among our nine additional putative hits (Supp. Tables 5, 6), only site 328 was identified by Radford and Bloom as conferring strong escape *in vitro* (escape score > 0.5). Many of these hits fall in the V4 variable region, where Radford and Bloom also found background-dependent escape mutations (e.g., locus 415 in BF520), although the two sets of loci did not overlap. Furthermore, several of Radford and Bloom’s strongest hits in other regions appearing in only one genetic background (sites 140, 151, 299, 327, & 330) were not observed *in vivo.* This suggests that, when extrapolating DMS results to intra-host settings, mutations durably appearing across DMS screening backgrounds are more likely to be relevant while background-dependent mutations may not.

We also examined changes in loop length and number of glycosylation sites in the variable loop regions. The loss of N332 glycan in V3 was the only change that was widely shared (10/10 participants without pre-existing high frequency resistance), but multiple other variable loop length and glycosylation patterns changed over time which might be linked to 10-1074 resistance (Supp. Fig. 6,7, Bai et al., 2024; Bricault et al., 2019; Zacharopoulou et al., 2024).

We therefore conclude that while loss of the N332 glycan and mutations at locus 325 appear to be the primary drivers of 10-1074 escape, other changes may also play a role.

### Low frequency preexisting resistance does not accelerate rebound timing or predict which variants dominate at rebound

Deeper sequencing revealed three 10-1074 participants with pre-existing escape motifs at N332 on day 0 (1HB3, 1HD4K, 1HD7K), which persisted through the study period (Fig. 2a, c, e). Surprisingly, pre-existing escape mutations were not associated with rebound timing or viral loads at weeks 0 or 1, after removing one outlier with pre-existing escape mutations at 100% frequency (Supp. Fig. 8). Furthermore, those pre-existing mutations did not dominate sequences recovered at later time points (for example, S334N in 1HD4K, Fig. 2b). A potential explanation for this pattern is that many escape mutations pre-exist below our detection limit or can easily be produced by mutation, such that escape mutations sampled at day 0 do not have an appreciable advantage over others. This may indicate that screening has limited clinical utility as a pre-treatment test for forecasting outgrowth of specific variants.

### Fitness differences among 10-1074 escape mutations

Because detectable pre-existing escape mutations did not dominate rebound populations, we tested if the amino acid abundances at sites 332 and 334 could be explained by nucleotide-specific mutation rates alone. N332T, N332I, and S334N were significantly enriched *in vivo* after 10-1074 administration compared to their relative probability of being generated by mutation (see Methods), suggesting greater *in vivo* fitness (Fig. 2e, Binomial tests, α=3.1×10^−3^𝑝=6.6×10^−9^,1.2×10^−4^,7.5×10^−35^ respectively). All amino acid substitutions at these sites equivalently remove the N332 glycan and confer 10-1074 resistance, so we tested if these apparent fitness differences could be associated with pleiotropic effects on other viral functions. Fitness differences were positively correlated with viral cell entry capabilities as measured by DMS (Site 332: Fig. 2f. Pearson 𝑟=0.6&𝑝=3.2×10^−5^, Site 334: Fig. 2g 𝑟=0.4 &𝑝=1.7×10^−2^, TRO11 clade-matched DMS backbone) and the favored mutation S334N was associated with a glycan shift in two participants (1HD4K & 1HD5K). This suggests viruses optimize secondary characteristics (e.g. cell entry) when escaping from 10-1074 *in vivo*.

### Distinct lineages within each participant acquire the same mutations for escape from 10-1074

The number of independent escape origins observed within each participant reflects how frequently escape mutations arise, an important consideration for combination therapy design (Coffin, 1995; LaMont et al., 2022). The presence of multiple distinct amino acid substitutions at sites 325, 332, and 334 within individual participants indicates that 10-1074 escape emerges through multiple independent origins. However, we predicted that deeper sequencing and linkage analysis may reveal additional cryptic resistance origins, in which distinct viral lineages acquire identical amino acid changes.

We first examined how often distinct nucleotide substitutions generated the same AA change following 10-1074 administration (i.e., both AAA and AAG encoding lysine), indicating multiple mutational origins. Among escape AAs with multiple possible codon encodings accessible from an ancestral AA via a single nucleotide change, the majority (14/24) were encoded in multiple ways within individual participants, indicating that more than one nucleotide substitution had occurred. Three distinct nucleotide substitutions could generate S334R, and one participant’s viral population, 1HD11K, contained all three of them (Fig. 3a). In contrast, no newly appearing AAs at loci other than 325, 332, or 334 were multiply encoded, even when reaching high frequencies.

**Figure 3.**
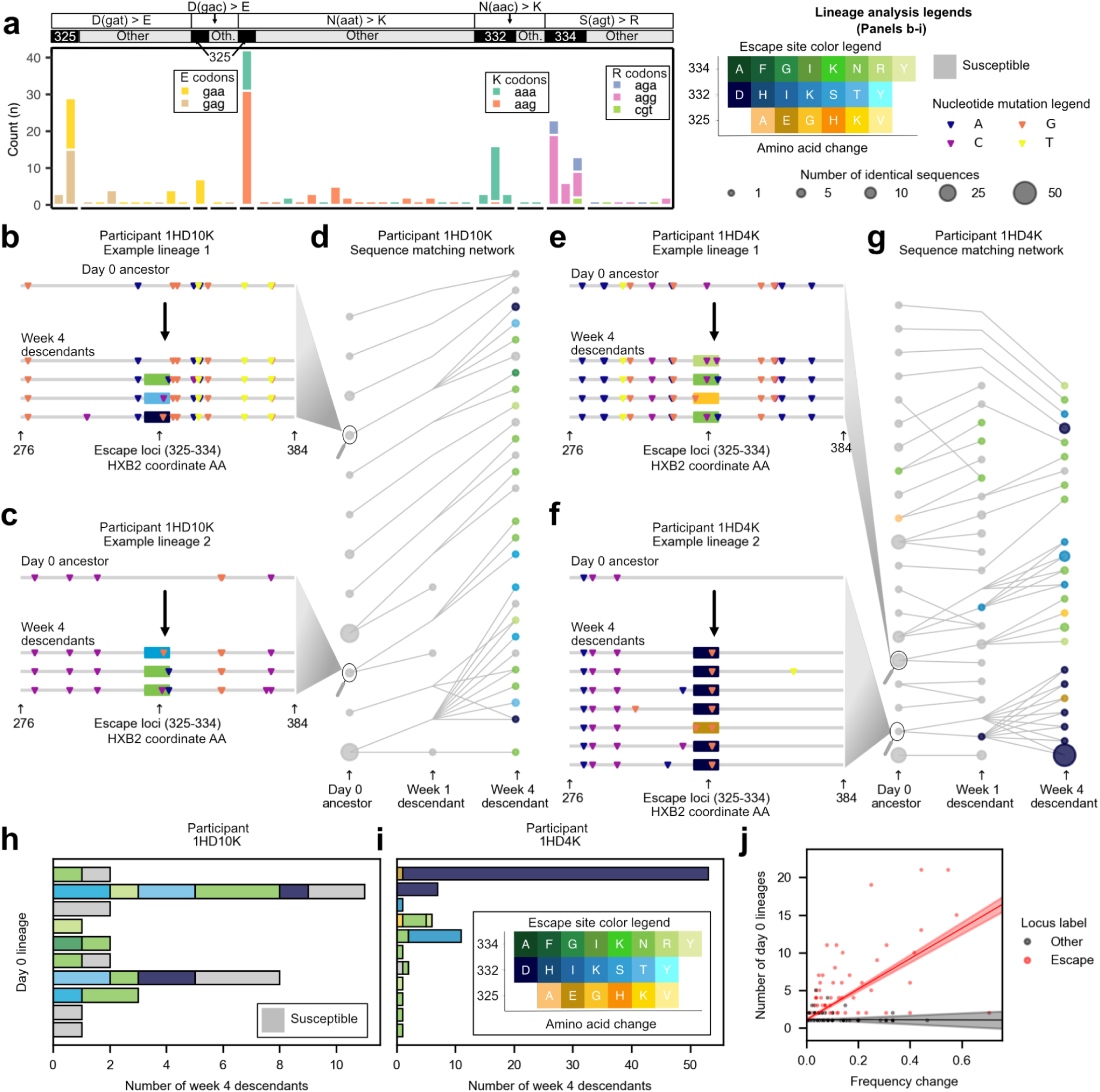
Escape from 10-1074 is highly parallel, with multiple occurrences of the same AA change spreading concurrently within participants. **(a)** Nucleotide codon encodings for AAs appearing after 10-1074 treatment stratified by ancestral AA nucleotide encoding. Each stacked bar represents nucleotide encodings for a single AA at a given locus from a single participant at an escape locus (325, 332, 334) or a non-escape locus (other). **(b-g)** Haplotype windows of 150bp flanking the 10-1074 resistance region were traced back to their common ancestors and matched via minimum Hamming distance in participants 1HD10K & 1HD4K. Highlighter plots (b,c,e,f) show haplotype examples of descendant-ancestor pairs. Network plots (d & g) indicate the descendant-ancestor relationships (edges) between sequences (nodes) at different timepoints. Sequences are connected to all equidistant ancestors. Sequences with multiple escape mutations are colored based on the escape mutation with the lowest HXB2 coordinate (applicable to panel f only). Haplotype networks for other participants are shown in Supplemental Figure 10 and are further summarized in Supplemental Table 7. **(h-i)** Bar plots of escape mutations carried by week 4 descendants of each day 0 lineage. Additional participants are shown in supplemental figure 11. **(j)** Number of day 0 lineages carrying each escape-associated (red) or non-escape-associated (grey) AA as a function of the AA’s maximum frequency increase during the trial. Escape-association was a significant predictor of the lineage count (Multiple linear regression, interaction coefficient for allele frequency and escape status: β_𝐴𝐴 𝐹𝑟𝑒𝑞𝑢𝑒𝑛𝑐𝑦:𝑁𝑜𝑛−𝑒𝑠𝑐𝑎𝑝𝑒_=− 20. 28, 𝑝 = 1. 03 × 10^−77^)

Because not all AAs that confer resistance can be multiply encoded, we further examined if the exact same escape mutation could be observed in tight genetic linkage to multiple viral backbones harboring different nearby mutations. We examined 150bp flanking either side of the escape motif (i.e., sites 325-334) and matched each unique post-treatment sequence to its closest ancestors at any previous time point (Fig. 3b-g, Supp. Fig. 10). By examining escape mutations in their lineage context, we found a median of 12 additional escape origins (defined as escape mutation-lineage combinations) beyond the original study (Supp. Table 7).

This analysis revealed two primary conclusions about parallel evolution in response to 10-1074. First, the same pre-treatment genetic background is often rescued by multiple distinct escape mutations (Fig. 3h,i, Supp. Fig. 11). Sequences which were identical at day 0 often appeared at later time points with different escape mutations, but no other changes in the flanking regions (for example, Fig. 3b,c,e,f). Second, the same escape mutations often emerge in parallel on distinct genetic backgrounds. We observed these patterns in all responding participants except 1HD7K, from whom only 6 sequences were recovered at week 4. We further found that escape mutations arose on significantly more genetic backgrounds than non-escape mutations (Fig. 3j, Multiple linear regression, interaction coefficient for allele frequency and escape status:

β_𝐴𝑙𝑙𝑒𝑙𝑒 𝐹𝑟𝑒𝑞𝑢𝑒𝑛𝑐𝑦:𝑁𝑜𝑛−𝑒𝑠𝑐𝑎𝑝𝑒_=− 20. 28, 𝑝 = 1. 03 × 10^−77^). Although rapid recombination can move escape mutations between genetic backgrounds, tightly conserved linkage structure on both sides of the escape motif suggest that at least some of these instances represent additional instances of independent mutational origins of therapeutic resistance that were previously undercounted by considering AA identity alone (Fig. 3b,c,e,f).

### Viral populations treated with 3BNC117 develop mutations at diverse loci that depend on sequence context

In contrast to 10-1074, no loci have been consistently identified as primary drivers of 3BNC117 escape (Bricault et al., 2019; Dingens et al., 2019; LaBranche et al., 2018; Radford & Bloom, 2025a; Scheid et al., 2016; Schommers et al., 2020). Consistent with this, we found no loci showing consistent AA changes between pre-and post-treatment timepoints across the majority of individuals receiving 3BNC117 (see Materials & Methods, Supp. Table 8), despite all study participants experiencing a dip and rebound in viral load.

To identify escape mutations without relying on convergent changes across participants and to advance beyond previous studies that report aggregate changes after bNAb infusion without identifying specific sites under selection, we conducted a selection scan with MPL, a method to deconvolute driving escape mutations from genetically linked passengers using longitudinal haplotype data (Sohail et al., 2021). MPL identified 11 loci under strong positive selection across seven different participants (Supp. Table 9, Supp. Fig. 12). These sites exhibited selection coefficients high enough (𝑠 ≥ 0. 044) to drive a substantial AA frequency increase during the trial (predicted to reach or exceed 15% frequency from a starting frequency of 5%, see Materials & Methods). In contrast to the convergent escape pathways observed for 10-1074, where participants developed similar mutations, responses to 3BNC117 were more heterogeneous, with only changes at locus 282 and in an insertion near locus 459 shared across two participants (Fig. 4a). Of our identified loci, 5/11 represent novel associations with 3BNC117 escape while the remaining 6/11 loci have either been previously linked to 3BNC117 escape (Bricault et al., 2019; Dingens et al., 2019; LaBranche et al., 2018; Radford & Bloom, 2025a; Scheid et al., 2016; Schommers et al., 2020) or fall in 3BNC117 contact sites (Schoofs et al., 2016; Zhou et al., 2013). In two participants, 2C1 and 2E1, MPL did not identify any loci under strong positive selection (Supp. Fig. 12). Note, running MPL considering sites as independent (i.e., ignoring linkage effects) yielded several additional hits highly linked (𝐷’ ≥ 0. 7) to putative escape mutations which we include in Supp. Table 10 as they may contribute epistatically to escape (“additional variation” in Fig 4a).

**Figure 4.**
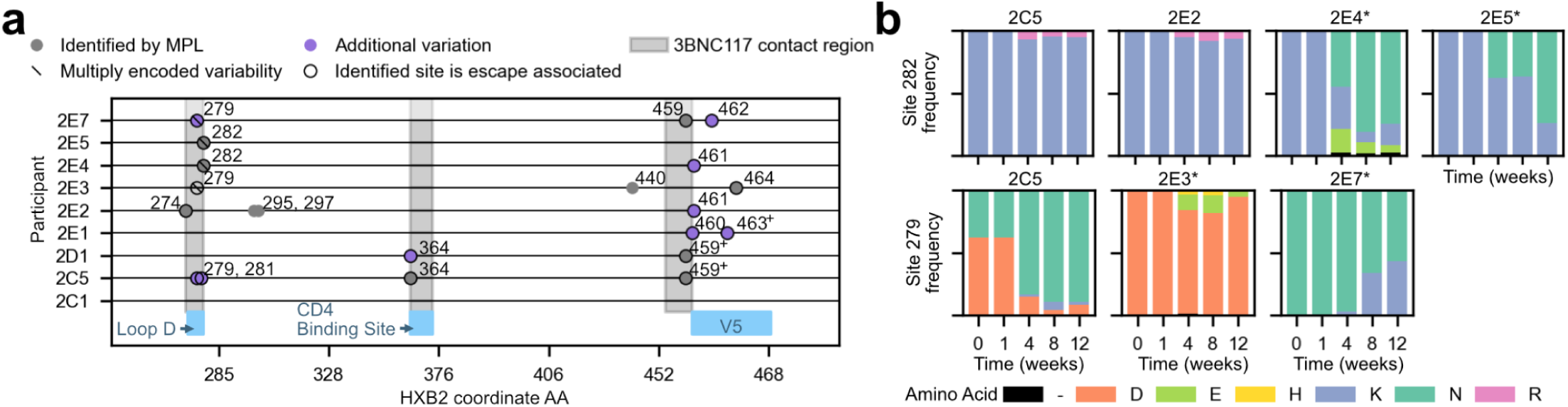
Loci changing following 3BNC117 administration exhibit only modest conservation across participants at the locus level and limited conservation at the AA level. **(a)** Putative escape loci against 3BNC117 identified in different participants (see Materials & Methods). Stacked symbols indicate that a locus was identified via multiple approaches. Black circular outlines denote loci with literature support (Bricault et al., 2019; Dingens et al., 2019; LaBranche et al., 2018; Radford & Bloom, 2025a; Schommers et al., 2020) or those falling in 3BNC117 contact regions (Schoofs et al., 2016; Zhou et al., 2013). Five loci without literature support outside of the contact regions (HXB2 coords 90, 97, 145, 706, and 754 in participants 2D1, 2E2, 2C5, 2E1, and 2E7, respectively) are not plotted (see supplemental tables 9 and 10 for all identified loci). ^+^denotes variation in an insertion relative to HXB2 at the listed coordinate. **(b)** Bar graphs of AA frequencies at escape loci identified in at least two participants. All participants with variation at each locus are shown, and * denotes participants where the locus was identified by a selection scan.

We hypothesized that if escape pathways were background specific, recurrent positively selected mutations may occur on genetically related backgrounds within a participant. We therefore repeated the multiple encoding analysis using data from the 3BNC117 cohort and found four additional loci with evidence for recurrent positive selection: locus 279 in two participants (2E3 and 2E7) and loci 97,145, & 706 in participants 2E2, 2C5, and 2E1, respectively (Fig. 4a). Additionally, mutations at MPL-identified locus 282 were multiply encoded in transmission pair members 2E4 and 2E5, corroborating their identification by MPL and supporting the hypothesis that related backgrounds may have more similar escape pathways. Of these loci, previous studies have linked mutations at 279 and 282 to 3BNC117 escape (Bricault et al., 2019; Schommers et al., 2020). Locus 97 has been linked to escape for other bNAbs targeting the CD4 binding site (Chen et al., 2021; Dingens et al., 2019; Huang et al., 2016a), but this is the first documentation of its importance in 3BNC117 escape.

In total, we identified loci under strong positive selection in 8/9 participants (excluding 2C1) (Supp. Tables 9, 10 and Figure 4a). Increased sequencing depth enabled more accurate measurement of AA frequency trajectories in key regions than previous analyses of these studies (Caskey et al., 2015; Schoofs et al., 2016) (Supp. Fig. 13-21). For example, in the original data, participant 2E3 had variation in loop D only 12 weeks after 3BNC117 infusion and at low frequency (D279E and D279H in 3/24 sequences, Caskey et al., 2015). However, deeper and more extensive sequencing revealed that these mutations circulated consistently from weeks 4 through 12 with a maximum frequency of 18% at week 8 (Supp. Fig. 18). Conversely, mutation 459E in participant 2D1 appeared fixed by week 4 in the original data, but deeper sequencing revealed that it remained below 27% frequency throughout the trial (Supp. Fig. 15). This highlights the potential for misinterpreting clinically-important *in vivo* allele frequency trajectories without sufficient sequencing depth.

In addition to clarifying the trajectories of previously identified mutations, we also found 11 AA identities in 5 participants at sites under strong positive selection which were not previously identified, (Supp. Table 11, Caskey et al., 2015; Schoofs et al., 2016). Many of these newly identified AAs (9/11) remained at low frequencies (<10%) but circulated at multiple timepoints. Together, these findings suggest that deeper sequencing enhances our ability to detect viral diversity and selection responses after bNAb infusion.

### Changes in V5 were associated with escape from 3BNC117

V5 is a known contact region for 3BNC117 and thus a likely site of viral escape. However, its high variability often undermines direct comparisons of specific loci across participants, a longstanding challenge in the field. We therefore examined non-single nucleotide variant based variation in V5 in the form of loop length and potential N-linked glycosylation (PNG) shifts and losses. In all participants, we found minimal changes in the V5 variable loop length and only two participants had minor decreases in the total number of V5 PNGs (2C5, 2E7, Supp. Fig. 22,23).

However, the location of V5 PNGs often shifted during the trial, displaying >10% frequency shifts in all participants except 2E3, 2E4 and 2E5 (Supp. Fig. 24, see Materials & Methods). These shifts occurred between PNGs at loci 460, 462, 463 and 465, although they did not show consistent directionality. Their magnitude was above 25% in participants 2C5, 2D1, 2E1, and 2E7 (Supp. Fig. 24). In 2C5, 2D1 and 2E7, these large shifts also corresponded to a single nucleotide variant trajectory identified by our selection scans as under positive selection. The heterogeneity of these findings across participants further suggests that specific motifs related to escape are dependent on the genetic context of the viral strain.

### Escape AA identities differ even across shared loci

We independently identified loci 279 and 282 as sites under selection in multiple participants, suggesting some sites of escape may be independent of background. However, which individual amino acid identities increased in frequency at those sites depended on pre-infusion genetic similarity (Fig. 4b). Both viral populations within the transmission pair of 2E4 and 2E5 acquired 282N repeatedly intra-host, but this mutation was not observed in any other viral population, suggesting it may be beneficial only in this particular genetic background. In contrast, we observed mutations at 279 exhibiting differing or even opposite responses in participants with genetically-distant viruses. Participants 2C5 and 2E7 experienced increases and decreases, respectively, in 279N frequency following 3BNC117 infusion. Meanwhile participant 2E3 experienced an increase in 279E, an AA identity not accessible by a single mutation for 2E7, indicating mutational accessibility may shape escape routes.

The dependence of escape mutations on genetic background builds upon a recent *in vitro* DMS experiment that found distinct 3BNC117 escape mutations in two highly divergent HIV genetic backgrounds (Radford & Bloom, 2025a). We found a lack of portability of escape mutations over a broader range of genetic distances (65-80% pairwise identity between strains here vs. 73% in Radford & Bloom), but identified convergent responses in highly related backgrounds (e.g. shared 282N response in the 96% identical transmission pair backgrounds, Fig. 4). We found almost no correspondence between the specific amino acids identified by Radford & Bloom and the *in vivo* trajectories (Supp. Fig. 25a), highlighting the limitation of DMS for cases of adaptation with high background dependency. At the locus level, loci shared across participants (279, 282) had higher escape scores than those loci we identified as under selection only in a single participant, although not significantly so (Supp. Fig. 25b). Loci mutated repeatedly across hosts *in vivo* may therefore represent escape routes that confer a fitness benefit across a greater range of genetic backgrounds, and may also be more readily identified by DMS.

### Intra-host 3BNC117 escape is often driven by parallel evolutionary responses

We examined how combinations of stringently filtered putative escape mutations (see Methods) expanded after 3BNC117 infusion to determine if multiple genetic backgrounds escaped in parallel within hosts, similar to viral responses to 10-1074. We discovered parallelism in escape responses in all participants except 2E1 and 2C1, who had minimal genetic change following 3BNC117 infusion (Fig. 5). In some participants, parallelism arose from multiple amino acid substitutions at a single locus (2C5, 2E4), whereas in others it involved substitutions at distinct loci with evidence of recombination between sites via the four gamete test (2E2, 2E7, 2E3) (Hudson & Kaplan, 1985). We also observed parallelism in escape responses in participants 2E3, 2E4, 2E5 and 2E7 as evidenced by escape AAs at sites 279 and 282 being encoded by multiple independent nucleotide changes, similar to patterns observed in 10-1074. However, unlike 10-1074, escape was confined to a limited number of genetic backgrounds, with a single dominant haplotype accounting for the majority of observed sequences.

**Figure 5.**
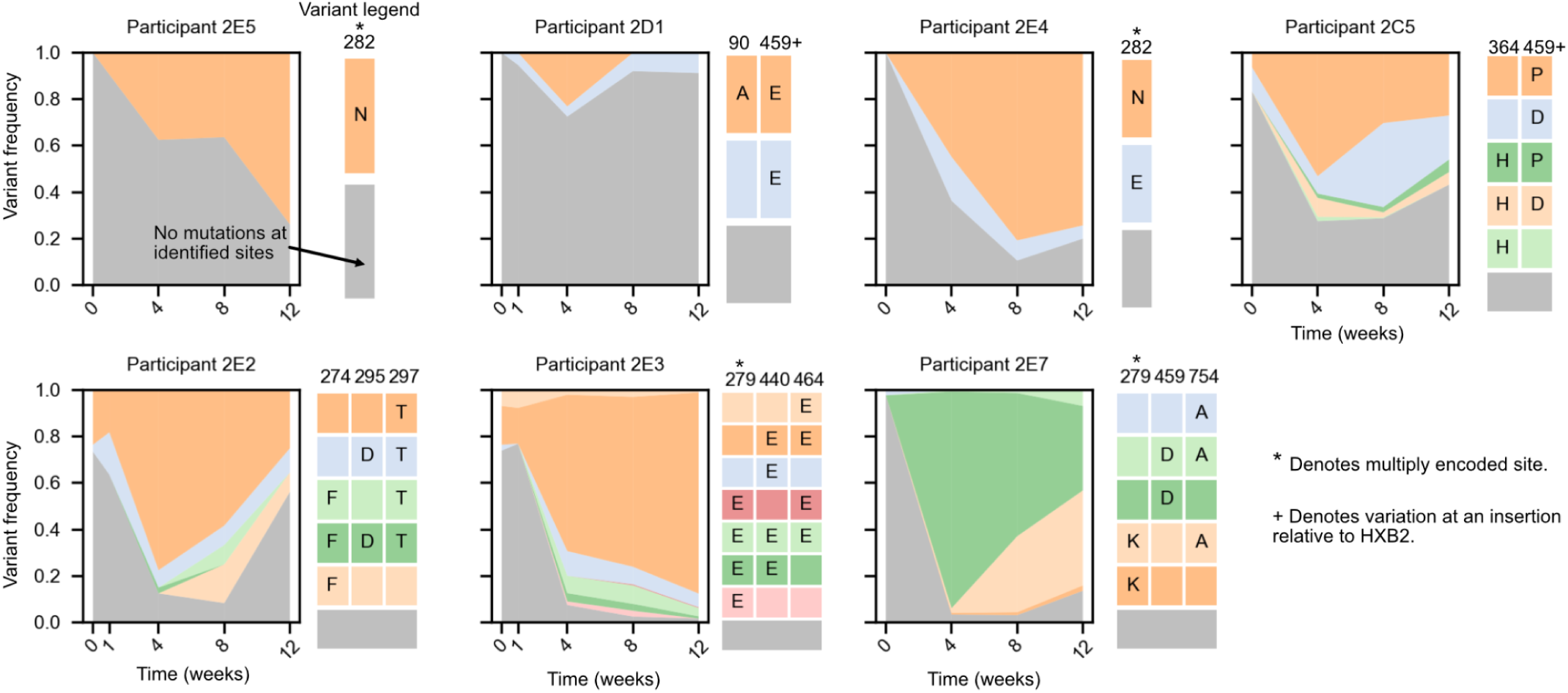
Parallel escape responses are observed in many 3BNC117-treated participants. **Left:** Frequencies of genetic backgrounds carrying positively selected loci identified by MPL or multiple encodings during the trial. Only loci with AAs increasing in frequency by >10% during the trial are shown. **Right:** AA combinations composing the genetic backgrounds plotted at left, matched by color and vertical ordering (mutations away from wild type shown). Starred (*) columns denote AAs with multiple observed nucleotide encodings and pluses (^+^) denote variation at an insertion site relative to HXB2. Sequences within the same variant group can differ at loci not identified by selection scans. Participants 2E1 and 2C1 carried no putative escape mutations exceeding 10% frequency and are not plotted.

This indicates that although multiple putative escape mutations arise concurrently in response to 3BNC117, they occur less frequently than in response to 10-1074.

## Discussion

Broadly neutralizing antibodies (bNAbs) are currently in development as both HIV prevention and cure modalities (Julg et al., 2022, 2024; Sneller et al., 2022; Williamson et al., 2025). The emergence of escape mutations in response to single bNAbs parallels early antiretroviral monotherapies (Larder et al., 1989), reinforcing the principle that durable suppression of HIV will likely require antibody combinations targeting multiple non-overlapping epitopes. In this study, we used deep haplotype-resolved sequencing to understand viral escape from bNAb monotherapy in people with chronic untreated HIV who received a single infusion of 10-1074 or 3BNC117.

Our study uses long-read SMRT-UMI sequencing to generate a large resource of nearly 7000 linkage-resolved HIV *env* sequences to track escape from two bNAbs targeting distinct epitopes. We achieve a 3.5-7.5x greater depth than prior studies and enable direct cross-trial comparison by applying the same modern approach to samples from two clinical studies. SMRT-UMI overcomes key barriers to *ex vivo* HIV sequencing, including limitations posed by *in vitro* recombination, abundant genetic diversity, and low frequency variation. Our analysis pipeline further provides a practical blueprint for investigating viral adaptive evolution in future trials.

With dense sampling and quantitative within-host evolutionary inference, our framework offers a sensitive and detailed longitudinal view of resistance emergence and viral dynamics under therapeutic pressure.

Collectively, these experimental and analytical innovations revealed new insights about viral escape from 10-1074 and 3BNC117, including: 1) previously unrecognized evolutionary parallelism in response to 10-1074 and 3BNC117, 2) *in vivo* evidence of background dependence in 3BNC117 responses as a function of genetic divergence between viruses, 3) new identification of putative escape mutations, especially in response to 3BNC117, 4) new evidence of low frequency preexisting resistance, and 5) apparent fitness differences between different 10-1074 escape mutations at loci 332 and 334. Importantly, because our analysis spanned two different bNAbs targeting distinct features of Env, we were able to directly compare how treatment-specific selective pressures shape resistance in fundamentally different ways.

HIV responses to 10-1074 were conserved across nearly all viral populations sequenced, but responses to 3BNC117 differed at the mutation level across participants. Specifically, in response to 10-1074, viral populations always mutated the GDIR motif at locus 325 or eliminated the N332 glycan to escape binding while viral responses to 3BNC117 involved a greater array of identified substitutions at the predicted binding sites.

These differences likely derive from the degree of viral conservation at the sites that interact with the two different bNAbs: 10-1074 contacts the N332 glycan and GDIR motif which are highly conserved (Sok et al., 2014), likely since small genotypic changes in this region can drive large phenotypic shifts such as tropism (Sok et al., 2016). 3BNC117 contact sites, particularly those in V5, vary substantially in comparison. As a result, different 3BNC117 escape mutations may be accessible in different populations, a feature that is shared with other bNAbs targeting this binding domain (Li et al., 2011). Our findings build on those from Radford and Bloom by showing that the strong background dependence they observed for *in vitro* 3BNC117 escape between clades A and B also operates *in vivo*, and arises intra-clade among clade B viruses (Radford & Bloom, 2025a). Our results therefore highlight a limitation of DMS in predicting *in vivo* trajectories where circulating variants diverge from laboratory strains. However, we do observe repeated 3BNC117 escape mutations intra-host and identical responses in a transmission pair who harbor closely related viruses, suggesting that accessible escape mutations may remain predictable among closely related strains (>95% sequence identity). The rapid emergence of background dependence implies that predicting even high frequency resistance from pre-treatment genetic screening may not be possible (as it is with ART) except in genetic backgrounds highly similar to those already profiled. Background dependence is also likely to adversely affect existing computational models that predict optimal combinations of bNAbs by assuming a fixed profile of escape mutations across backgrounds (LaMont et al., 2022).

An important question is how our observations for 10-1074 and 3BNC117 extend to other antibodies that target the same epitopes. V3-glycan bNAbs engage a comparatively constrained epitope; V3 length is relatively consistent across circulating viruses and the glycan scaffold is often conserved, which likely contributes to the higher predictability we observe for escape from V3 glycan binding antibodies (Mouquet et al., 2012; Stephenson et al., 2021). In contrast, while the CD4 binding site itself is also conserved, CD4 binding site targeting bNAbs have variable footprints and modes of access to bypass Loop D and/or V5, creating more opportunity for background-dependent selection as we see across participants treated with 3BNC117 (Huang et al., 2016b; Li et al., 2011; Schommers et al., 2020). These epitope-level differences provide a mechanistic basis for why 10-1074 and 3BNC117 differed in our study and inform how escape may generalize across bNAbs within each class.

We also found major differences in the degree of parallelism between 10-1074 and 3BNC117 escape. By identifying recurrent AA substitutions, we discovered that 10-1074 escape mutations arise from a median of 22 origins within a single population. This greatly exceeds HIV responses to 3BNC117 observed here and to early antiretroviral therapies (Feder et al., 2016, 2021; Pennings et al., 2014; Williams & Pennings, 2020), which are typically driven by one to three origins of resistance. Greater parallelism in viral escape from 10-1074 compared to 3BNC117 likely reflects a larger number of accessible escape mutations (Hermisson & Pennings, 2017). Because escape from 10-1074 appears more parallel than escape from early ART, it is possible that the combinatorial challenge necessary for certain bNAbs combinations might be even higher than for ART. It is also possible that intra-host genetic responses for 10-1074 and early ART are similarly parallel and this study’s sequencing depth, the genetic diversity of *env* versus *pol,* and identity of escape mutations (i.e., multiple identifiably-distinct AA substitutions conferring escape) allow greater discovery of parallelism for 10-1074 than ART. For 3BNC117, despite the lesser degree of parallelism, discovering repeated production of escape mutations suggests this antibody, and other CD4-binding site antibodies, may face similar obstacles.

Despite their differences, both 10-1074 and 3BNC117 harbored rare pre-existing resistance mutations at day 0 (3/11 harbored detectable resistance mutations to 10-1074 and 2/9 harbored putative resistance mutations to 3BNC117). Identifying these mutations was potentiated by deeper sequencing, but any attempt to comprehensively sequence a viral population *ex vivo* will be incomplete. While this appears to present an obstacle for predicting viral responses to bNAb therapy, surprisingly, we found that rare pre-treatment resistance mutations for 10-1074 did not predict which mutations reached highest frequencies after treatment. In agreement with our parallelism results, this finding suggests that large viral populations can readily produce escape mutations to individual bNAbs and that attempting to screen for rare pre-existing mutations may have limited clinical applications.

Profiling viral responses *ex vivo* further allowed us to understand HIV bNAb escape in the presence of the other intra-host selection pressures. For example, we found viral populations favor 10-1074 escape mutations that preserve cell entry functionality or shift compensatory glycans. Evasion of a person’s endogenous HIV antibodies or cellular mediated immunity via CD8+ T cells or NK cells is likely to further constrain *in vivo* escape pathways (Borrow et al., 1997; Chung et al., 2011; Elemans et al., 2017; Gao & Barton, 2025), which may further contribute to the diversity of 3BNC117 responses within this cohort. Escape mutations reverting back to pre-treatment AAs in participants from both trials (Fig. 2A-D, Fig 5) indicate that the fitness costs of escape from either bNAb are sufficiently strong to revert once the virus is not under active selection from either 10-1074 or 3BNC117. Measuring these *in vivo* dynamical phenomena represents another important reason to investigate escape in clinical cohorts in addition to via *in vitro* approaches like DMS.

We note three important caveats for this study. First, we cannot unequivocally establish neutralization or escape beyond previously published neutralization data. For 10-1074, escape mutations at loci 325, 332, and 334 have been well documented as conferring complete resistance to 10-1074 (Bricault et al., 2019; Caskey et al., 2017; Dingens et al., 2019; Radford & Bloom, 2025; Selzer et al., 2025). For 3BNC117, we cross referenced our findings with existing DMS and neutralization studies, but we can only classify variants as escape-associated, and we cannot strongly assert that any single mutation causes escape from 3BNC117.

This is a challenge shared with other *in silico* escape prediction approaches (Bai et al., 2019; Magaret et al., 2019; Moldt et al., 2021). Second, we leverage linkage patterns to identify 3BNC117 escape mutations but discriminating between neutral and selective dynamics is inherently challenging, even when equipped with comprehensive time series data. Rapid changes in viral population sizes can result in large allele frequency fluctuations even under neutral conditions (Swan et al., 2022). This challenge is exacerbated when viral backbones carry multiple putative escape mutations rising in frequency together, which could result from synergy between mutations or genetic hitchhiking. We account for this by conservatively using a strong selection coefficient cut-off in evaluating our MPL results to limit false positives and using corroborating information about multiple nucleotide encodings and shared responses across participants when available. Third, we note that our sequencing was based solely on circulating virus from plasma. Although it is possible that viral resistance dynamics may differ in tissue specific sites, the blood plasma compartment has been shown to be relatively representative of the viral population within tissues during viremia, mitigating the impact of compartmentalization on our results (Chaillon et al., 2020; Kearney et al., 2015; Pardons et al., 2025).

Several trials have now combined bNAbs targeting two or three non-overlapping epitopes either as primary therapy or after an ART interruption (Bar-On et al., 2018; Gaebler et al., 2022; Julg et al., 2024; Mendoza et al., 2018; Sneller et al., 2022). These combinations can achieve sustained periods of ART-free remission in certain people living with HIV, but some viral populations nevertheless rebound prematurely, especially those treated with two bNAb therapies. Adding additional components like long-acting ART drug lenacapavir (Eron et al., 2024), or VRC07.523LS and cabotegravir (Taiwo et al., 2025) or an immune-modulation TLR9 agonist (Peluso et al., 2025) have achieved even greater success. Furthermore, bNAbs potentially offer functions beyond viral suppression, such as engaging and possibly stimulating the host immune response, and when the virus is highly sensitive they may achieve outcomes that rival or exceed ART (Fumagalli et al., 2025; Lu et al., 2016; Niessl et al., 2020; Rosás-Umbert et al., 2022; Schoofs et al., 2016; Tebas, 2025). Deep sequencing of resistance profiles to determine how HIV evades these antibodies lays an important foundation for the rational design of next-generation antibody combinations and adjunctive strategies to achieve durable, ART-free remission.

## Methods

### Sample selection

All participants were provided informed consent and the parental studies (Caskey et al., 2015, 2017) were approved by the Rockefeller University IRB.

#### MCA-0885

Plasma samples were selected from 12 participants in a phase 1 clinical trial testing a single 10-1074 broadly neutralizing antibody (bNAb) infusion in viremic participants living with HIV-1 (NCT 02511990, Caskey et al., 2017). Baseline and subsequent timepoints were sequenced. (Fig. 1b, Supp. Table 1).

#### MCA-0835

Plasma samples were selected from 7 participants in a phase 1 clinical trial testing a single 3BNC117 bNAb infusion in viremic participants living with HIV-1 (NCT 02018510, Caskey et al., 2015). Baseline and subsequent timepoints were sequenced (Fig. 1b, Supp Table 2).

### RNA isolation and cDNA synthesis

Frozen plasma samples (1mL) were thawed and viral RNA isolated using MinElute Virus Spin kits (Qiagen, #55704) or Viral RNA Mini spin kits (Qiagen, #52904). cDNA synthesis performed from viral RNA using HIV-1 specific oligos or SMRT-UMI cDNA 5’ ultramers containing random barcodes (Westfall et al., 2024). cDNA was synthesized using SuperScript IV Reverse Transcriptase (10U/uL, Thermo Fisher Scientific), and purified using RNAClean XP beads (Beckman Coulter) at a 1:1 ratio.

### Amplification

HIV *env* cDNA concentration was estimated using an end point dilution as previously described (Westfall et al., 2024). Samples with low viral load, where cDNA was synthesized without SMRT-UMI oligos, were amplified at limiting dilution, to ensure positive PCRs contained only a single cDNA template. PCR 2 primers contained 5’ tag sequences complementary to unique index primers added during a third PCR to allow for pooling of positive PCRs for sequencing.

For SMRT-UMI sequencing, a maximum of 50 HIV *env* cDNA molecules were input into each 25uL PCR reaction. PCR products were pooled and cleaned by magnetic bead size selection (AMPure XP, 0.5x) and a second-round PCR was performed using 1uL of bead cleaned 1_st_ round product. A heteroduplex resolution step was conducted at the end of the second PCR, which consisted of spiking in additional enzyme, 2_nd_ round forward and reverse primers and dNTPs to each reaction tube. Final amplified products were confirmed by a 3kb band visualized by gel electrophoresis and purified by magnetic bead size selection (AMPure XP magnetic beads, 0.7x).

Specific details for these steps can be found in the supplement along with all cDNA and PCR primers (**Supp. Methods, Supp. Table 13)**. Index primers are listed along with the demultiplexing pipeline at https://github.com/lcohnlab/sga-pipeline.

### Library Prep

PCR product was quantified using the Qubit HS 1x dsDNA assay (ThermoFisher). Up to 16 SMRT-UMI sample IDs or 384 individually indexed PCR were pooled in equimolar amounts in a ratio equivalent to the estimated number of templates input into that sample’s PCR. Only one timepoint from each participant was included in each pool to avoid incorrect identification of samples as contamination by bioinformatic processing.Libraries were prepared using the SMRTBell Express Template Prep kit 2.0 (Pacific Biosciences). Sample pools were divided into 90ng input reactions (10uL each) and barcoded using the Barcoded Overhang Adaptor Kit - 8A (Pacific Biosciences). Libraries were sequenced using a PacBio Sequel IIe (Fred Hutchinson Cancer Center Genomics and Bioinformatics Core Facility).

### Sequence generation and sequence preparation

Demultiplexing and consensus generation was performed from PacBio CCS reads separately from single positive PCR or samples amplified in bulk with SMRT-UMI oligos. Reads from single positive PCRs used a simple pipeline (https://github.com/lcohnlab/sga-pipeline), while the PORPIDpipeline software was adapted to run on a local cluster for samples amplified in bulk. PORPIDpipeline performs read quality filtering, read demultiplexing by sample and UMI, exclusion of erroneous UMI, consensus sequence generation for each UMI, contamination check, sequence quality filtering, and report generation as previously described (Westfall et al., 2024). Additional sequence filtering took place for each SMRT-UMI sample using custom R scripts. First, the sequences were ordered from those with the most reads contributing to their consensus to those with the least. Next, the sequences consisting of the bottom 15% or 25% of total reads were excluded, depending on which cutoff excluded the lowest tail of the read distribution best. Adapted PORPIDpipeline software and additional filtering scripts available at https://github.com/lcohnlab/PORPIDPipeline_Cluster-AC-2025.

High-quality consensus sequences from PORPID for each participant were trimmed to matching autologous *env* sequences (Caskey et al., 2015, 2017; Schoofs et al., 2016) and aligned to HXB2 reference genome *env* sequence using MUSCLE 5.1 (AliView Alignment Viewer, Larsson, 2014). Sequences with premature stop codons or frameshift mutations following polynucleotide sequences of 4 or more identical nucleotides were frameshift corrected. Nucleotide sequences were translated to corresponding amino acid sequences and re-aligned to the HXB2 reference *env* amino acid sequence. We excluded sequences with internal deletions totaling more than 3% of the *env* sequence as well as insertions greater than 3% of the sequence.

### Sequencing validation

To determine whether recombination occurs during reverse transcription, we combined RNA from two infectious molecular clones, 89.6 (Collman et al., 1992) and LAI.2 (Wain-Hobson et al., 1985), for cDNA synthesis, performed single genome amplification (Palmer et al., 2005; Keele et al., 2008) on the resulting cDNA, and sequenced single viral envelope genes by Illumina MiSeq. Compared to the reference 89.6 and LAI.2 genomes, no 89.6/LAI.2 recombinants were identified, demonstrating a negligible amount of recombination during cDNA synthesis (Supp. Fig. 26a).

We tested the ability of our adapted PORPID pipeline ((Westfall et al., 2024), adaptations in methods) to filter out PCR recombinants during bulk amplification by combining 89.6 and LAI.2 cDNA, which contained different sample ID barcodes, processing them with bulk PCR amplification, and sequencing them by PacBio.

Compared with the reference 89.6 and LAI.2 envelope sequences, we observed no recombinant sequences identified following PORPID filtering (Supp. Fig. 26b).

We further confirmed that our approach and the original single genome amplification (SGA) recovered similar viral populations at the same time point using a pairwise panmixia test (Caskey et al., 2015, 2017; Kearney et al., 2014; Schoofs et al., 2016, FDR-adjusted p < 1 x 10^-4^). Of the samples with matched SGA sequences, 34/43 had no significant compartmentalization between sequencing methods. Of those that were significantly compartmentalized, the inadequate sample size of SGA or the presence of a subclade found by SMRT-UMI but not by SGA contributed to the compartmentalization (Supp. Fig. 2).

## Code availability

All additional code for conducting analyses listed below is available at https://github.com/evromero-uw/hiv_bnabs_paper_archive.git.

### Sequence alignment and phylogenetic analysis

Within host *env* nucleotide sequences were codon-aligned using MUSCLE v5 and then manually inspected and edited using AliView. SMRT-UMI sequences, recovered here, were aligned along with available full *env* SGA sequences from NCT 02511990 participants collected in Caskey et al., 2017 and from NCT 02018510 participants collected in Schoofs et al., 2016 & Caskey et al., 2015. Phylogenetic trees were created using IqTree2 (Minh et al., 2020) and plotted using the R packages ape and phytools (Paradis & Schliep, 2019; Revell, 2024). All HXB2 positions reported are relative to the start of the *env* gene as outlined by the LANL HIV database (*Numbering Positions in HIV Relative to HXB2CG*, n.d.).

### Identification of potential escape loci

We analyzed alignment positions with <10% gaps and a day 0 majority allele frequency ≥50%. For each participant and position, we identified the day 0 majority amino acid (AA) and calculated the proportion of sequences differing from this AA at subsequent timepoints. We report sites with majority AA loss >0.4 (10-1074 cohort) or >0.05 (3BNC117 cohort). No sites were variable in most 3BNC117 participants, prompting an additional selection scan.

We ran an additional selection scan using MPL with run parameters matching those used by Sohail et al., 2021 (mutation rates from Zanini et al., 2017,𝑁_𝑒_ = 1 × 10^4^, gaussian regularization strength 𝑔 = 5 × 10^4^, <95% alignment column gap frequency, >3 sequences/time point). Analyses included data through week 8 to correspond with therapeutic bNAb serum concentrations. Sequences for participant 2D1 were truncated at HXB2 nucleotide position 1530 due to missing trial day 0 sequencing data. We chose a filtering cutoff of 𝑠 ≥ 0. 044 to identify sites where positive selection could create a substantial frequency increase during the trial (reaching or exceeding 15% frequency from 5% over 8 weeks or 28 generations under a drift-free, discrete generation model of haploid reproduction, Felsenstein, 2019, p.58).

For 3BNC117, we ran a single locus variant of MPL that ignores linkage (off-diagonal entries of the covariance matrix are set to 0, but all other parameters remain the same). Sites in 3BNC117 contact regions with s≥0.044 in the single-locus but not full model were classified as “additional variation” and excluded unless noted.

We determined an escape locus had literature support if it 1) had a significant escape score, as determined by dms_tools2 findSigSel, against the relevant bNAb in a recent DMS experiment (Bloom, 2015; Radford & Bloom, 2025a), or was reported as escape-associated in 2) the LANL HIV antibody contact database (Bricault et al., 2019; Dingens et al., 2019; LaBranche et al., 2018; Los Alamos National Laboratory, n.d.; Schommers et al., 2020) or the LANL CATNAP IC50 data by the antibody database program (West et al., 2013; Yoon et al., 2015, run parameters: min residue freq = 1%, max # of rules= 10, free set = 20%). We denote sites identified by the above methods that also had literature support as “putative escape loci”.

### Variable loop characterization

Potential N-linked glycosylation sites (PNGs) were identified via the canonical NX[ST] amino acid motif, where X is any amino acid excluding proline (Bause & Hettkamp, 1979). Any participant where the majority PNG pattern at day 0 changed in frequency by 10% or more during the trial was denoted as having shifts in glycosylation patterns.

### Analysis of 10-1074 mutational probabilitie**s**

We computed the relative probabilities of mutating away from N332 or S334 via all possible single step nucleotide substitutions based on participant-specific codons present at day 0 and HIV-specific mutation rate estimates (Zanini et al., 2017). Codons for escape mutations present at day 0 were excluded from the analysis. Expected relative frequencies of escape AAs at sites 332 and 334 in each participant were then calculated by weighting the mutation rate-adjusted accessible escape AAs for each ancestral codon by its participant-specific day 0 frequency. Enrichment as plotted in figure 2 was calculated as the log ratio of observed to expected AA frequency in each participant. To test the statistical support for enrichment, we tested if the proportion of escape sequences at a given site with a specific AA identity was higher than its expectation using a one-sided binomial test with Bonferroni correction, after pooling the observed counts and taking a sample size weighted average of the expected proportions across participants.

### Multiply encoded variant analysis

We identified all AA identities within a participant sampled only after day 0 and determined if multiple distinct nucleotide changes from codon triplets present at day 0 could produce the AA substitution. We restricted this analysis to newly appearing AAs to minimize the impact of secondary synonymous substitutions. Sites were reported as multiply-encoded if at least two sequences carried two or more different nucleotide encodings, and singly encoded otherwise. Sites in poorly aligned regions were excluded.

### Ancestor network analysis

We traced haplotypes carrying 10-1074 escape mutations in a 300bp window flanking the GDIR motif and 332 glycan (HXB2 nucleotide coordinates 825-1152 with sites 975-1002 masked). This window size was chosen to minimize the impact of viral recombination during the initial four weeks of therapeutic response given existing estimates of HIV generation time and recombination rate (Perelson et al., 1996; Romero & Feder, 2024), but our results were qualitatively robust to window size. Each sequence sampled at weeks 1 and 4 was assigned to its closest ancestor at any previous time point via minimum hamming distance. Sequences equidistant from multiple ancestors were assigned to all possible ancestors. The haplotype network graph was formed with each node representing a unique 300 bp motif and each edge connecting a descendant motif to its closest matched ancestor. Each connected component of the network represents a single lineage deriving from day 0 and consists of sequence motifs more closely related to each other than to any others sampled. We defined the number of origins of 10-1074 escape within each participant as the number of unique escape AA and lineage combinations within the participant.

We tested if 10-1074 escape alleles appeared on more day 0 sequence motifs than non-escape alleles, controlling for allele frequency. We repeated the network analysis described above in which we centered 300bp windows on all sites of non-escape variation and determined the number of day 0 lineages on which non-escape alleles appeared. We compared escape and non-escape alleles by fitting the following linear regression:

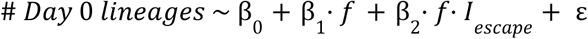

in which f represents the maximum frequency of an allele at any point during the trial, and 𝐼_𝑒𝑠𝑐𝑎𝑝𝑒_ is an indicator variable of whether or not the allele is associated with escape. We report the strength, directionality and significance of the interaction coefficient, β_2_.

### 3BNC117 haplotype analysis

We report haplotypes composed of all alleles at putative escape loci that were not the day 0 majority and increased ≥10% during the trial, supported by ≥2 sequencing reads at the timepoint.

### Comparison to deep mutational scanning data

Published DMS data for TRO11 and BF520 Env were filtered to include variants observed ≥2 times (TRO11) or ≥3 times (BF520), per author recommendations (Radford & Bloom, 2025b). For amino acid–level analyses, mean escape scores across DMS replicates were compared to frequency changes of matched amino acids at the same HXB2 coordinates in vivo. For locus-level analyses, we used the maximum mean escape score across all amino acids at the locus and both strains.

## Acknowledgements

We thank all of the study participants who participated in the first-in-human trials of 10-1074 and 3BNC117; the Rockefeller University Hospital nursing staff and Clinical Research Support Office and nursing staff; Deysi Dubon for laboratory support, Anthony West, Daniel Reeves, John P. Barton, Pleuni S. Pennings, and Michel Nussenzweig for critical reading of our manuscript; all members of the Feder and Cohn labs, Alexis Chang, Caelan Radford, and Jesse Bloom for helpful discussion; Wenjie Deng for bioinformatics support. This work was supported in part by the following NIH NIAID grants UM1AI64565 and U01AI169385 to LBC and MC, the Pew Research Scholars Program to LBC, and NIH 1DP2CA280623-01 and the Gilead Sciences Research Scholars Program in HIV to AFF; This work was additionally supported by the Genomics and Bioinformatics Shared Resource (Research Resource Identifier [RRID]:SCR_022606) of the Fred Hutchinson Cancer Center/University of Washington/Seattle Children’s Cancer Consortium funded by the National Cancer Institute (grant number P30 CA01570).

