## Supplemental Figures for "Recurrent mutations drive rapid HIV escape from two broadly neutralizing antibodies *in vivo*"

**a**

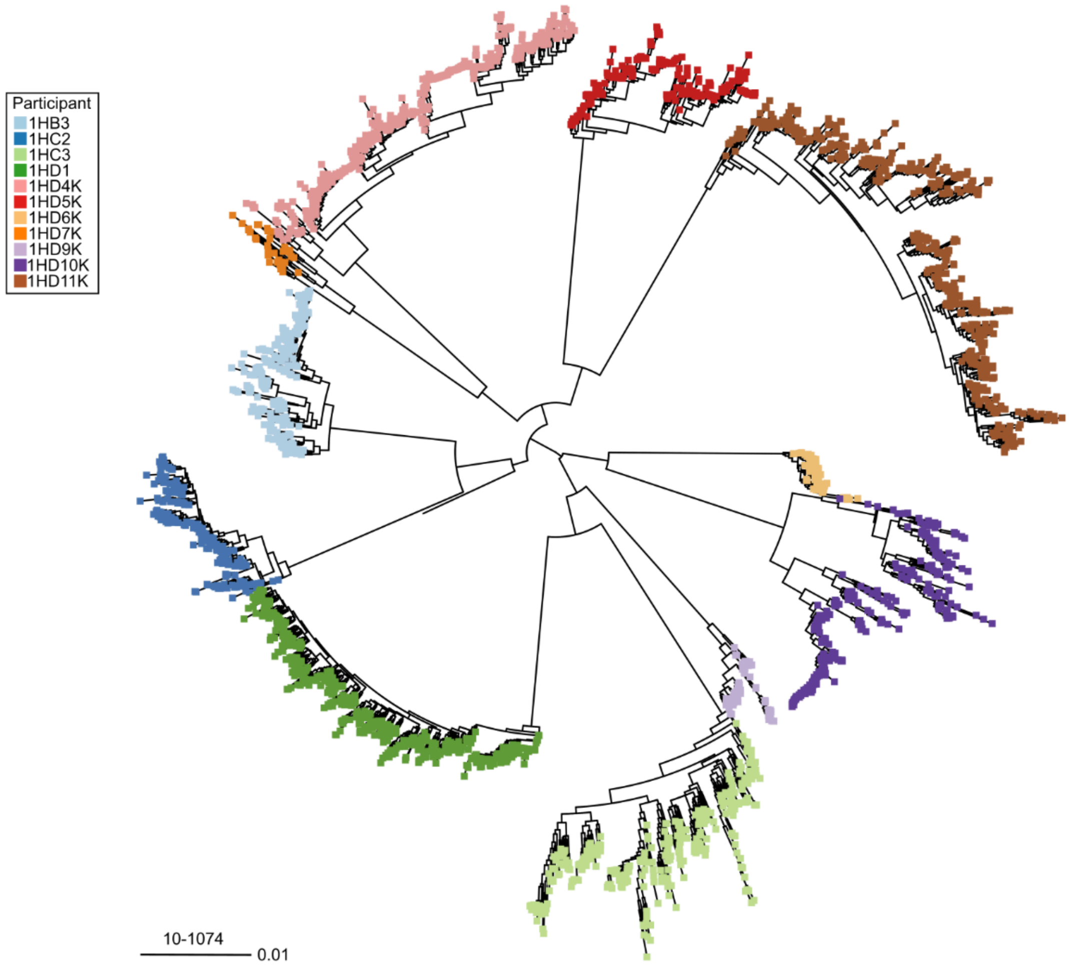
Participant
1HB3
1HC2
1HC3
1HD1
1HD4K
1HD5K
1HD6K
1HD7K
1HD9K
1HD10K
1HD11K
10-1074
0.01

**b**

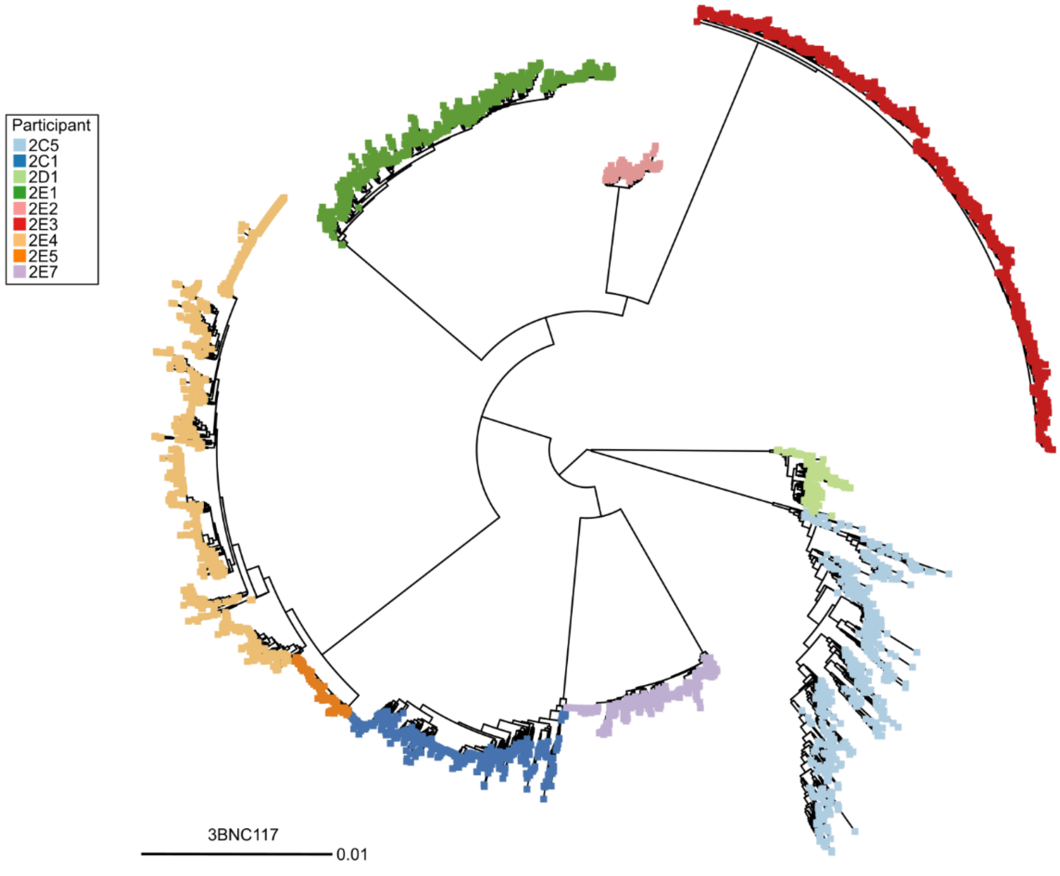
Participant
2C5
2C1
2D1
2E1
2E2
2E3
2E4
2E5
2E7
3BNC117
0.01

**Supplemental Figure 1:** Intermixed tree of 10-1074 **(a)** and 3BNC117 **(b)** sequences clustering by participant. SMRT-UMI *env* sequences and previously published *env* sequences from NCT 02511990 and NCT 02018510 were plotted using IQTree2 and the Tamura-Nei substitution model (Caskey et al., 2015, 2017; Minh et al., 2020; Schoofs et al., 2016). Sequences clustered by participant as expected. Participants 2E4 and 2E5 in the 3BNC117 group show less genetic divergence from each other and were previously reported to be a transmission pair (Caskey et al, 2016).

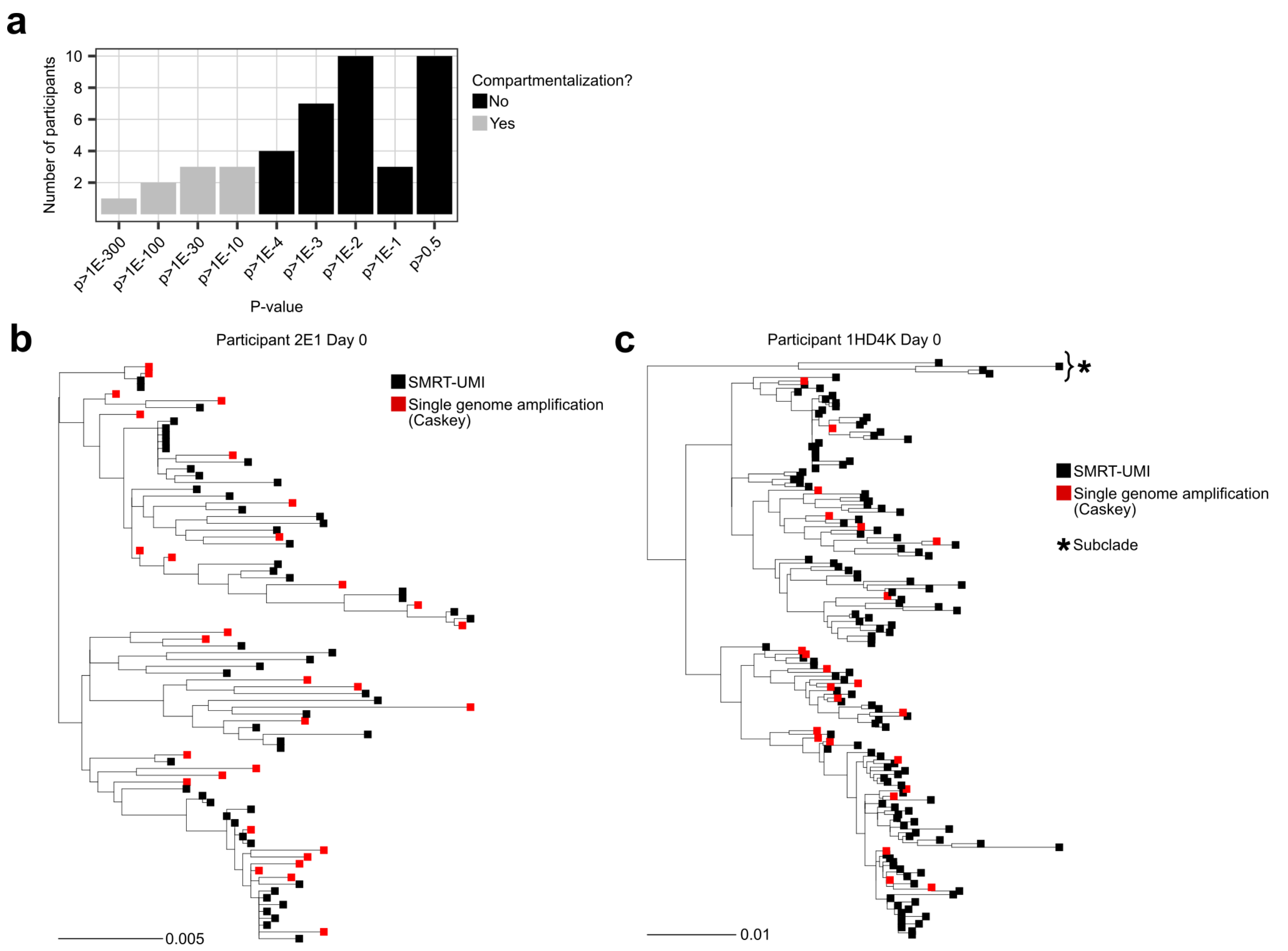


**Supplemental Figure 2:** Assessment of compartmentalization and panmixia of Env sequences using the single genome amplification method and the SMRT-UMI method.

**(a)** Validated SMRT-UMI sequences were compared to sequences originally recovered using single genome amplification (SGA) from the same samples to determine if the two sequencing approaches recovered similar viral populations using a pairwise panmixia test (Caskey et al., 2015, 2017; Kearney et al., 2014; Schoofs et al., 2016). Of the samples with matched SGA sequences, 34/43 were found to have no significant compartmentalization between sequencing methods ($p < 10^{-4}$, see Materials & Methods). **(b)** Example tree of a matched sample (2E1 Day 0) showing panmixia between SMRT-UMI and SGA sequences ($p = 0.07$) **(c)** Example tree of a matched sample (1HD4K Day 0) showing significant compartmentalization between SGA and SMRT-UMI sequences ($p < 3 \cdot 10^{-4}$). In this example, the presence of a subclade found by SMRT-UMI (*) but not SGA contributed to compartmentalization. Subclade existence was further confirmed by its sampling in both methods at later timepoints. All samples that exhibited compartmentalization with $p < 10^{-4}$ had either inadequate SGA or SMRT-UMI sequence recovery or the presence of an unsampled subclade.

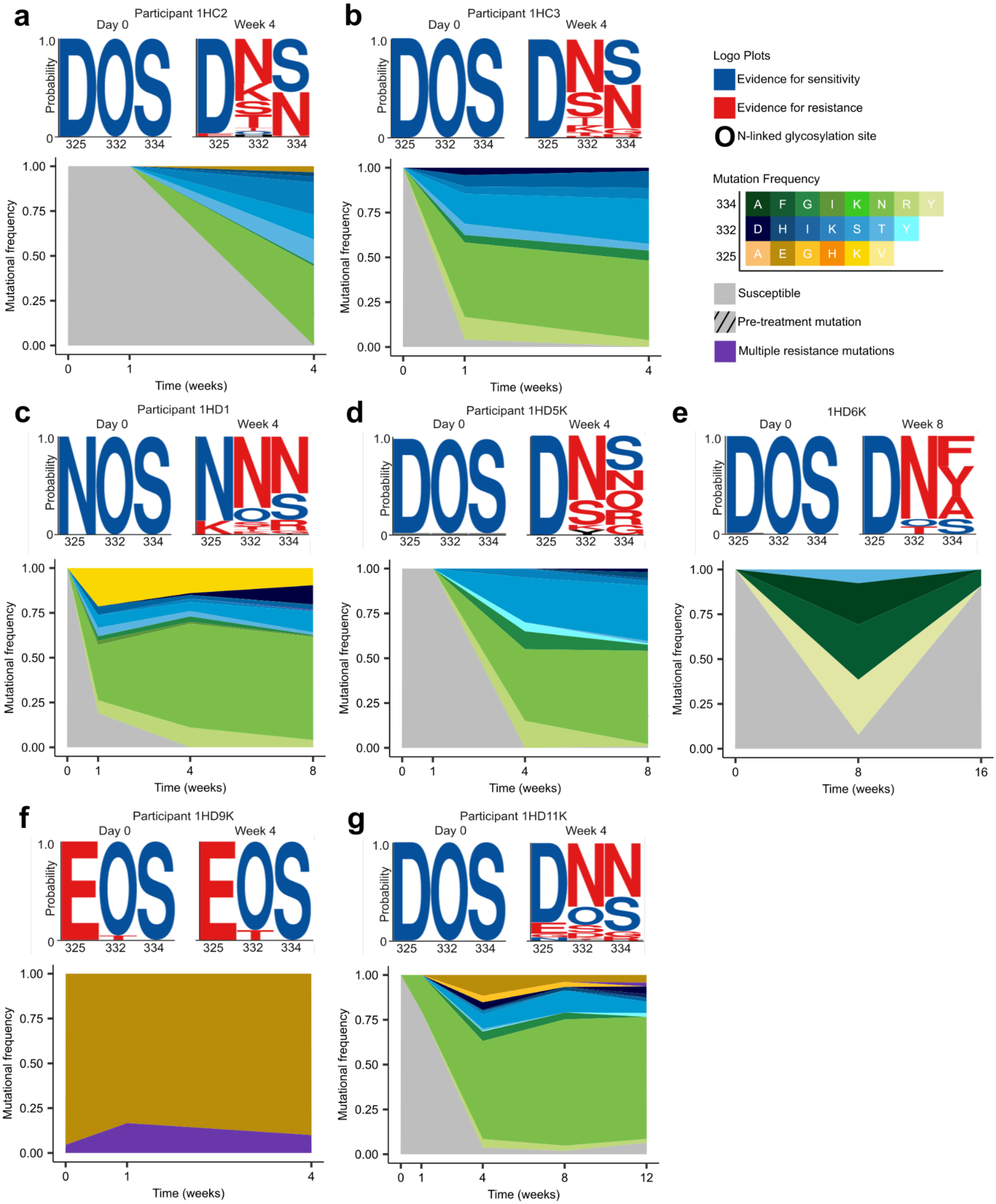

a
Participant 1HC2
Day 0
Week 4
b
Participant 1HC3
c
Participant 1HD1
d
Participant 1HD5K
e
1HD6K
Week 8
f
Participant 1HD9K
g
Participant 1HD11K
Probability
Mutational frequency
Time (weeks)
325
332
334
Logo Plots
Evidence for sensitivity
Evidence for resistance
N-linked glycosylation site
Mutation Frequency
334 A F G I K N R Y
332 D H I K S T Y
325 A E G H K V
Susceptible
Pre-treatment mutation
Multiple resistance mutations

**Supplemental Figure 3.** Cohort-wide dynamics of viral escape mutations following 10-1074 treatment. **(a-g)** (top) Logo plots of amino acids at loci 325, 332, and 334 are shown for pre-treatment time point (day 0) and the first post-nadir time point (week 8 in 1HD6K, week 4 in all others) in additional participants treated with 10-1074. AAs in red or blue have previously been reported to confer resistance or sensitivity to 10-1074, respectively (Bricault et al., 2019; Caskey et al., 2017; Dingens et al., 2019; Radford & Bloom, 2025). (bottom) Frequency plots of escape mutations over time for each participant, with each mutation indicated by color and hatching indicating those escape mutations discovered pre-treatment.

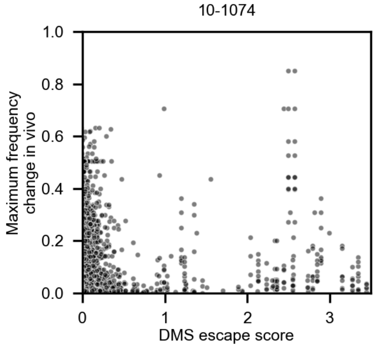


**Supplemental Figure 4**. Deep mutational scanning (DMS) escape scores against 10-1074 for Env AAs on both TRO11 and BF520 backgrounds (Radford & Bloom, 2025) compared to their *in vivo* maximum frequency change in any 10-1074-treated participant during the trial. Higher escape scores indicate less antibody neutralization by 10-1074. Loci with no equivalent site in the DMS envelopes are omitted.

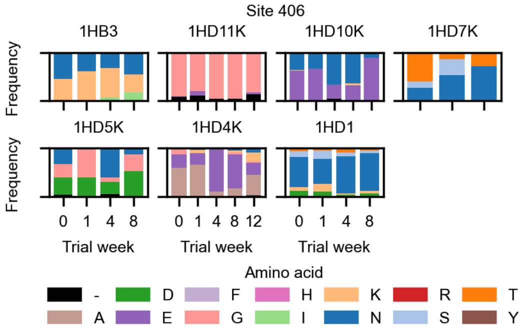


**Supplemental Figure 5**. Allele frequencies at HXB2 site 406 in 10-1074-treated participants over time. Each subpanel shows the AA frequencies for a single participant and participants with any variation are shown.

10-1074

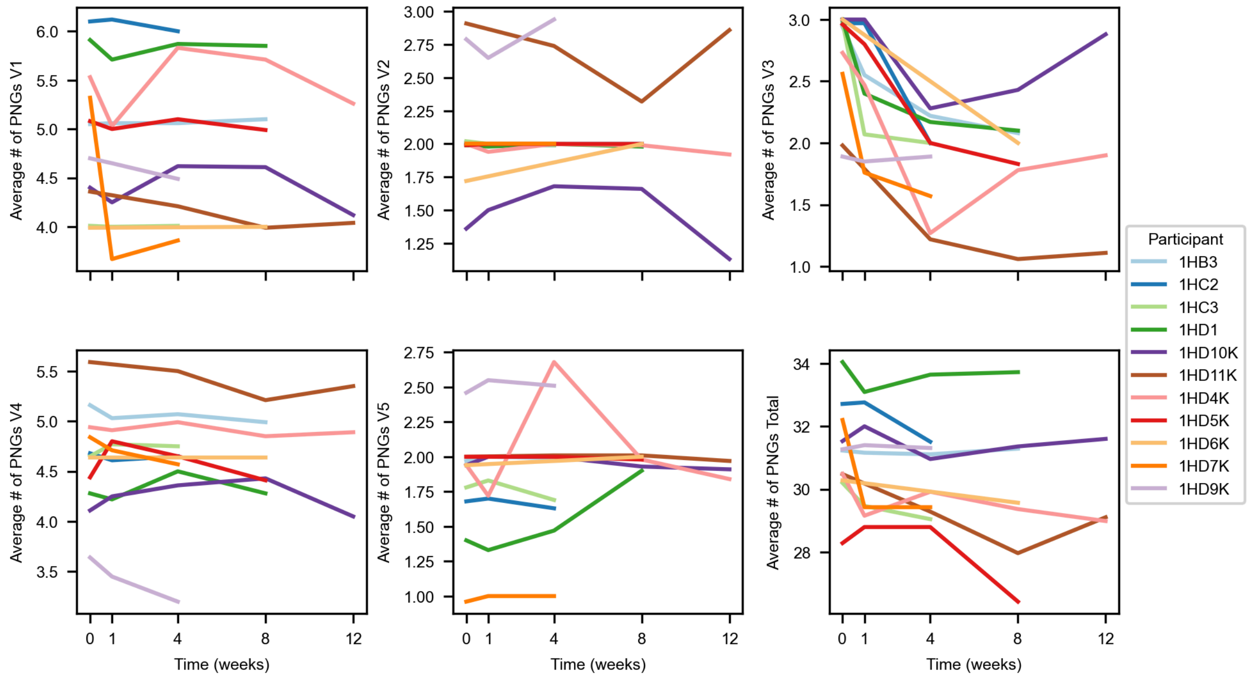


**Supplemental Figure 6.** Average numbers of potential N-linked glycosylation sites (PNGs) within each participant throughout the trial in the 10-1074-treated cohort. Each panel displays data for a single variable loop (V1-V5) or the total count of PNGs summed across all variable loops.

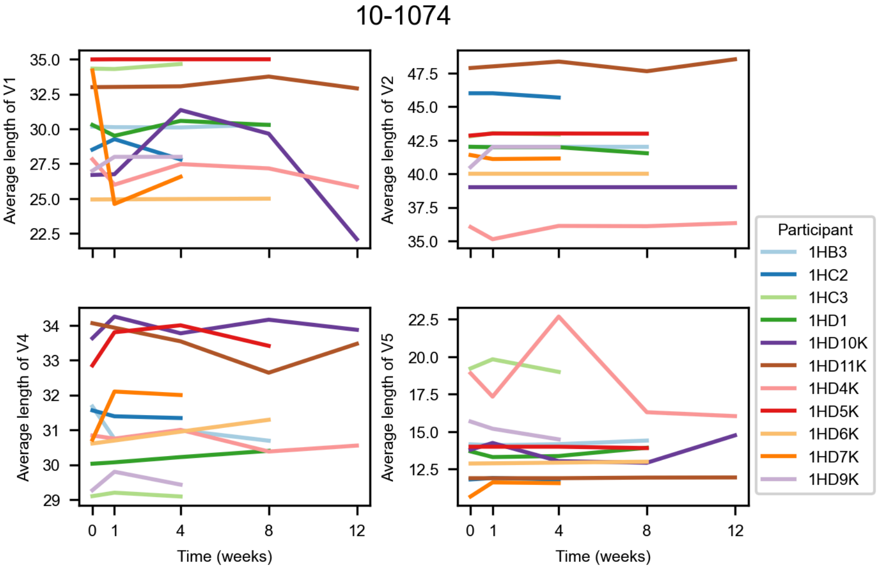


**Supplemental Figure 7.** Average lengths of variable loop regions throughout the trial in participants treated with 10-1074. V3 is not shown as its length is highly conserved (De Wolf et al., 1994; Laakso et al., 2007; Zhong et al., 1995).

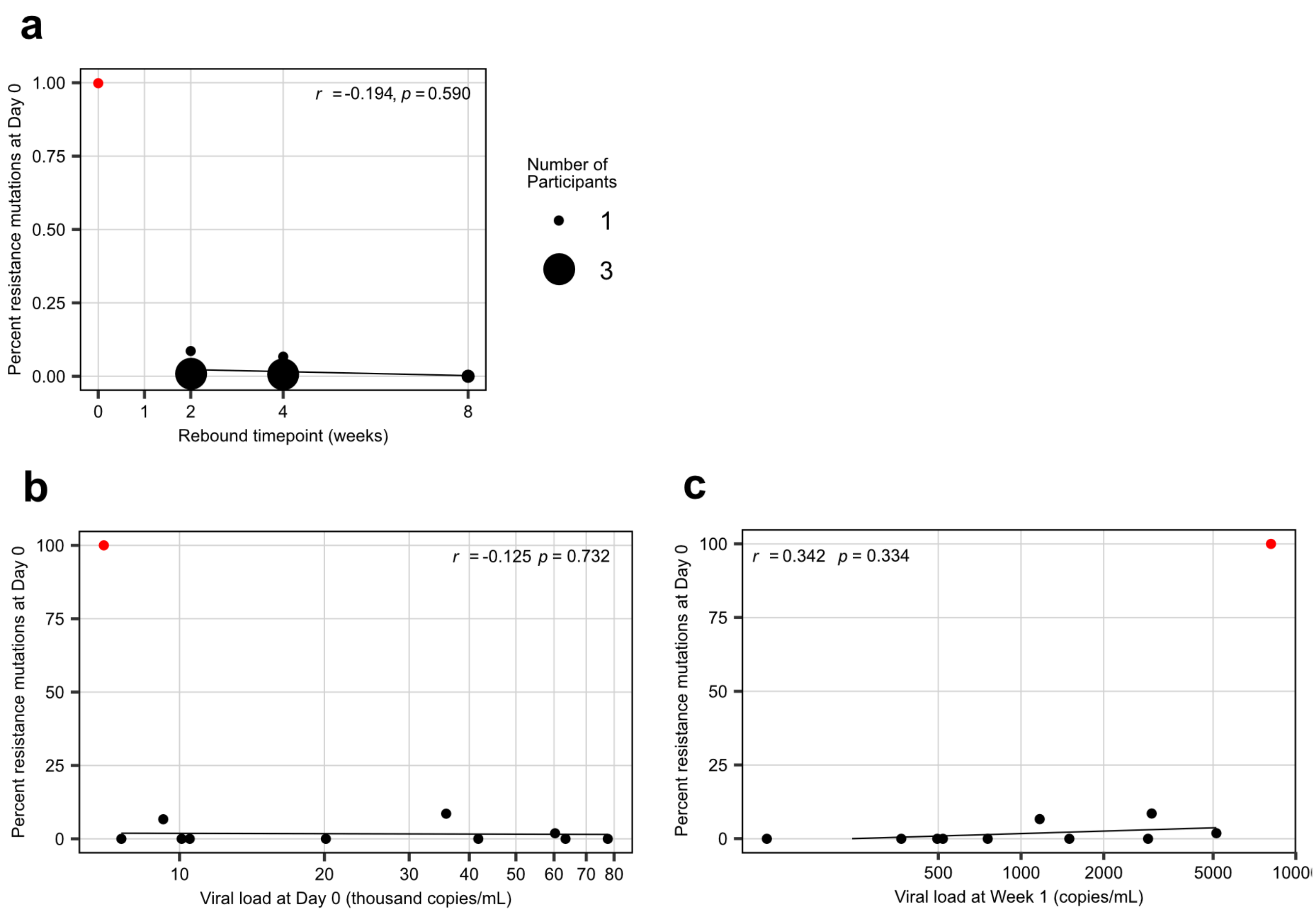


**Supplemental Figure 8:** Pre-existing 10-1074 resistance does not correlate with viral load or faster viral rebound when excluding one participant with a resistance mutation at 100% frequency at day 0 (1HD9K, plotted in red). **(a)** Pre-existing resistance mutations do not correlate with faster viral rebound after 10-1074 administration (Pearson $r = -0.914$, $p = 0.590$). The rebound timepoint was defined as the first timepoint post-nadir with an increase in viral load of $0.5\ log_{10}$copies/ml or greater, and subsequently confirmed at the next timepoint. **(b)** Pre-existing resistance mutations did not correlate with viral load at day 0 (Pearson $r = -0.125$, $p = 0.732$). **(c)** Pre-existing resistance mutations did not correlate with viral load at week 1 timepoint (Pearson $r = 0.342$, $p = 0.334$).

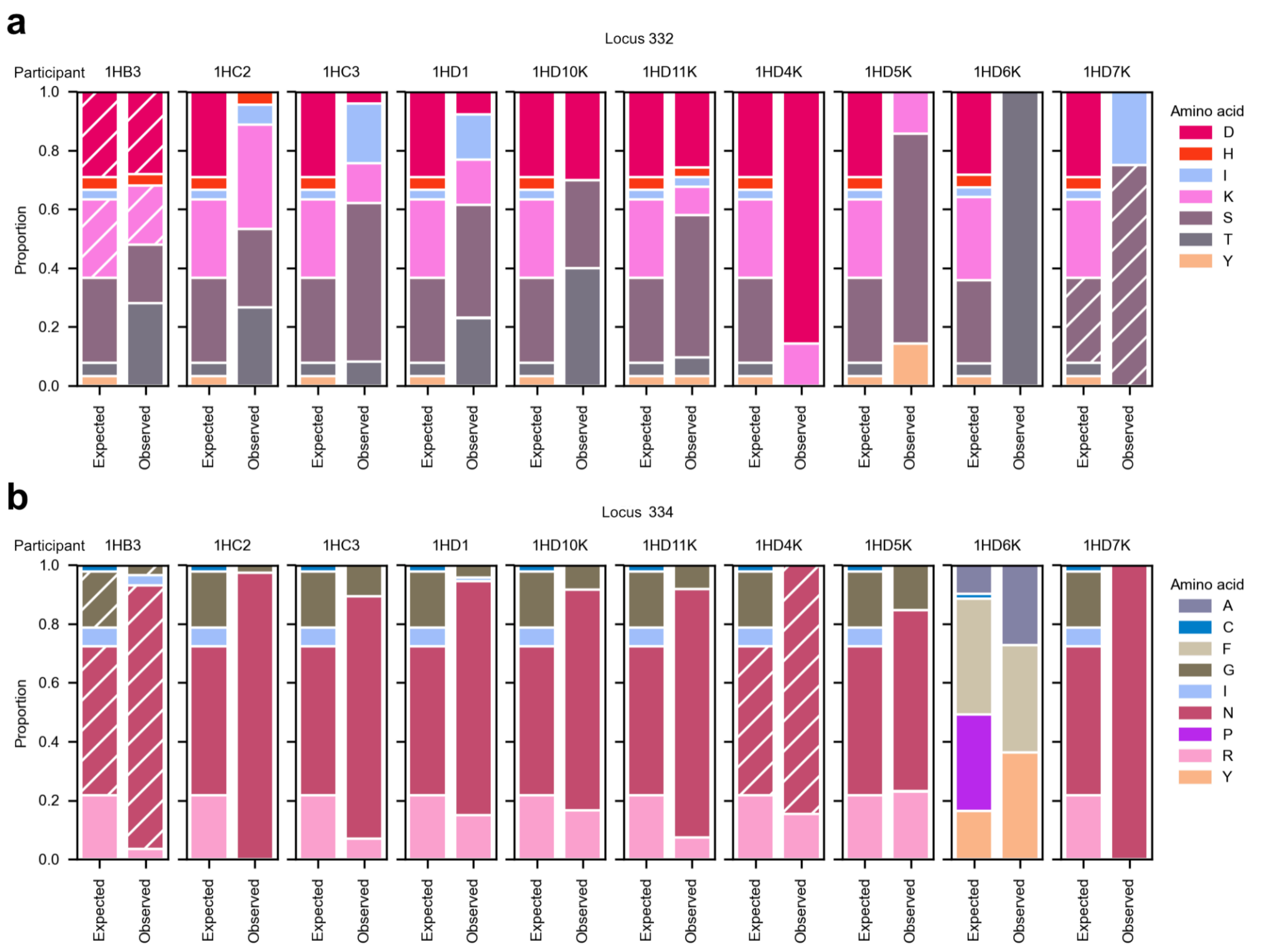


**Supplemental Figure 9.** Enrichment of specific AAs associated with 10-1074 escape. **(a,b)** Bar plots of the expected and observed proportions of escape mutations at escape loci 332 & 334 at trial week 4. Expected AA distributions were calculated using ancestral codons observed at day 0 and published single nucleotide mutation rates (Zanini et al., 2017, see Materials & Methods). Hatched texture indicates AAs sampled at day 0 that predated 10-1074 treatment.

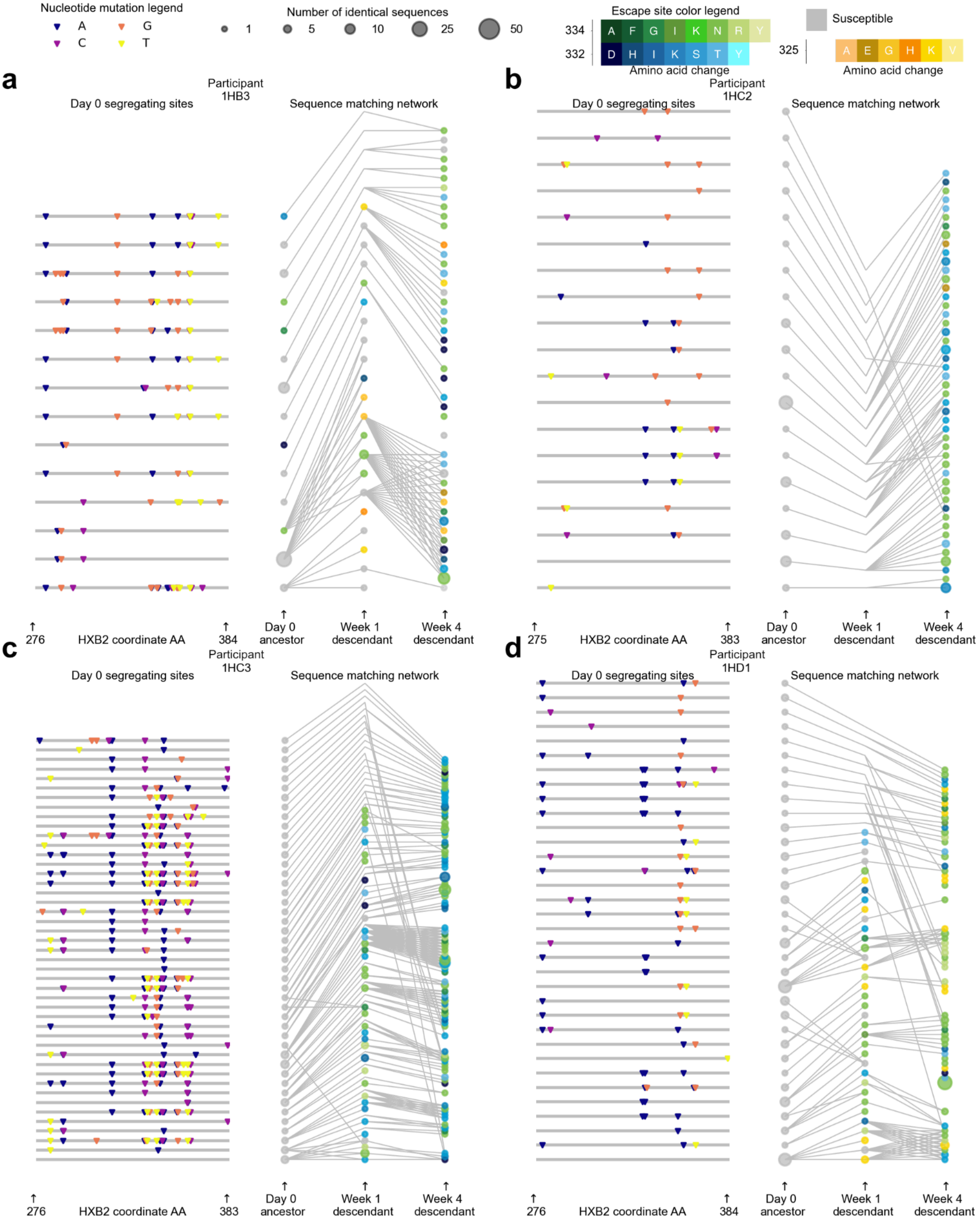

Nucleotide mutation legend
A
G
C
T
Number of identical sequences
1
5
10
25
50
Escape site color legend
334
A F G I K N R Y
332
D H I K S T Y
Amino acid change
Susceptible
325
A E G H K V
Amino acid change
a
Participant 1HB3
Day 0 segregating sites
Sequence matching network
276
HXB2 coordinate AA
384
Day 0 ancestor
Week 1 descendant
Week 4 descendant
b
Participant 1HC2
Day 0 segregating sites
Sequence matching network
275
HXB2 coordinate AA
383
Day 0 ancestor
Week 1 descendant
Week 4 descendant
c
Participant 1HC3
Day 0 segregating sites
Sequence matching network
276
HXB2 coordinate AA
383
Day 0 ancestor
Week 1 descendant
Week 4 descendant
d
Participant 1HD1
Day 0 segregating sites
Sequence matching network
276
HXB2 coordinate AA
384
Day 0 ancestor
Week 1 descendant
Week 4 descendant

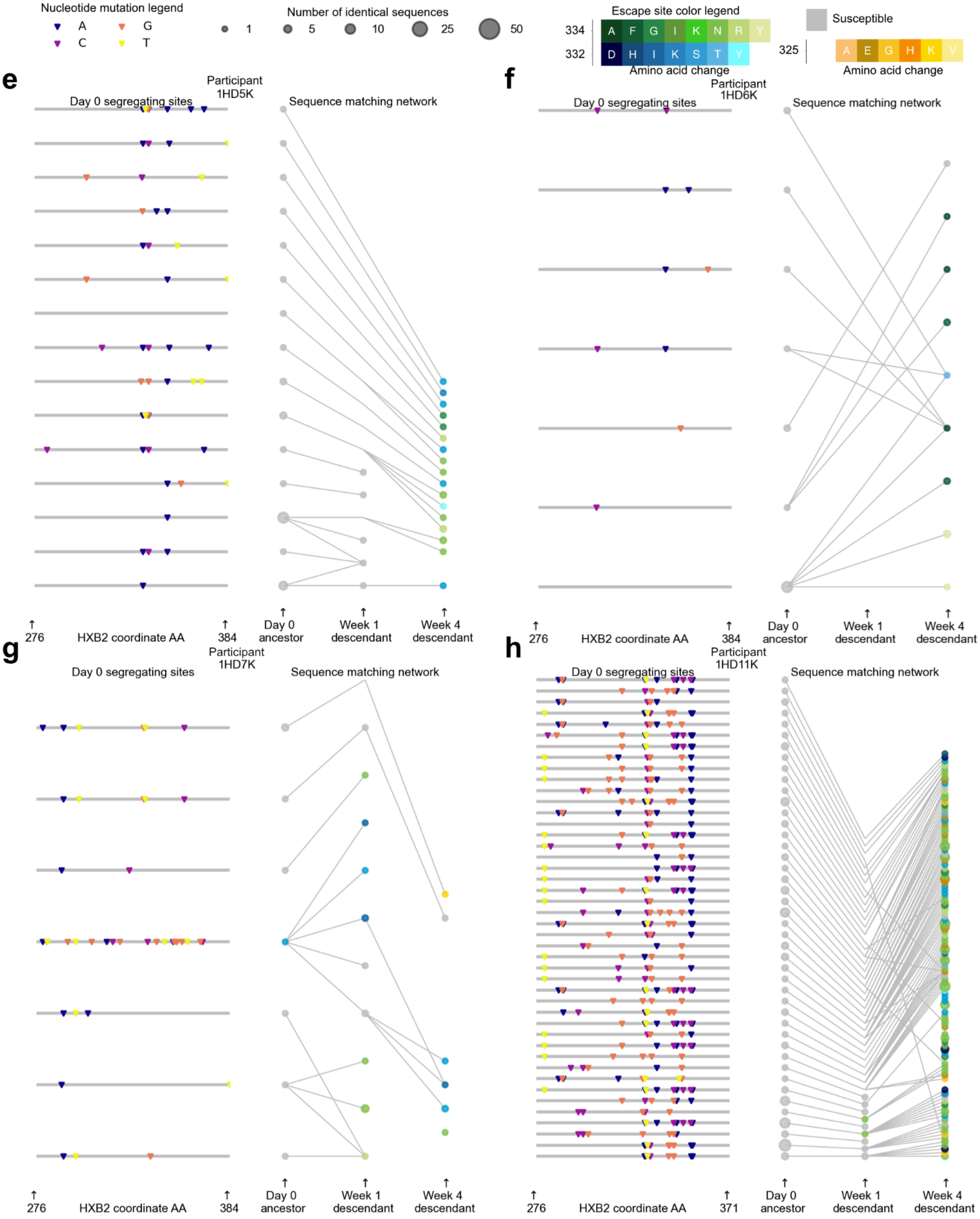

Nucleotide mutation legend
A
G
C
T
Number of identical sequences
1
5
10
25
50
Escape site color legend
334
A F G I K N R Y
332
D H I K S T Y
Amino acid change
Susceptible
325
A E G H K V
Amino acid change
e
Participant 1HD5K
Day 0 segregating sites
Sequence matching network
276
HXB2 coordinate AA
384
Day 0 ancestor
Week 1 descendant
Week 4 descendant
f
Participant 1HD6K
Day 0 segregating sites
Sequence matching network
276
HXB2 coordinate AA
384
Day 0 ancestor
Week 1 descendant
Week 4 descendant
g
Participant 1HD7K
Day 0 segregating sites
Sequence matching network
276
HXB2 coordinate AA
384
Day 0 ancestor
Week 1 descendant
Week 4 descendant
h
Participant 1HD11K
Day 0 segregating sites
Sequence matching network
276
HXB2 coordinate AA
371
Day 0 ancestor
Week 1 descendant
Week 4 descendant

**Supplemental Figure 10.** All 10-1074 treatment responders show evidence of multiple lineages escaping 10-1074 in parallel. **(a-h)** Haplotype windows of 150bp flanking the 10-1074 resistance region were traced back to their common ancestors and matched via minimum Hamming distance in participants 1HB3, 1HC2, 1HC3, 1HD1, 1HD5K, 1HD6K, 1HD7K & 1HD11K. Day 0 highlighter plots (left subpanels) indicate the haplotype of the corresponding (y-axis position matched) day 0 ancestor in the right subpanel. Network plots (right subpanels) indicate the ancestor-descendant relationships (edges) between sequences (nodes) at different timepoints. Sequences are connected to all equidistant ancestors. Other participant networks are available in figure 3b-g.

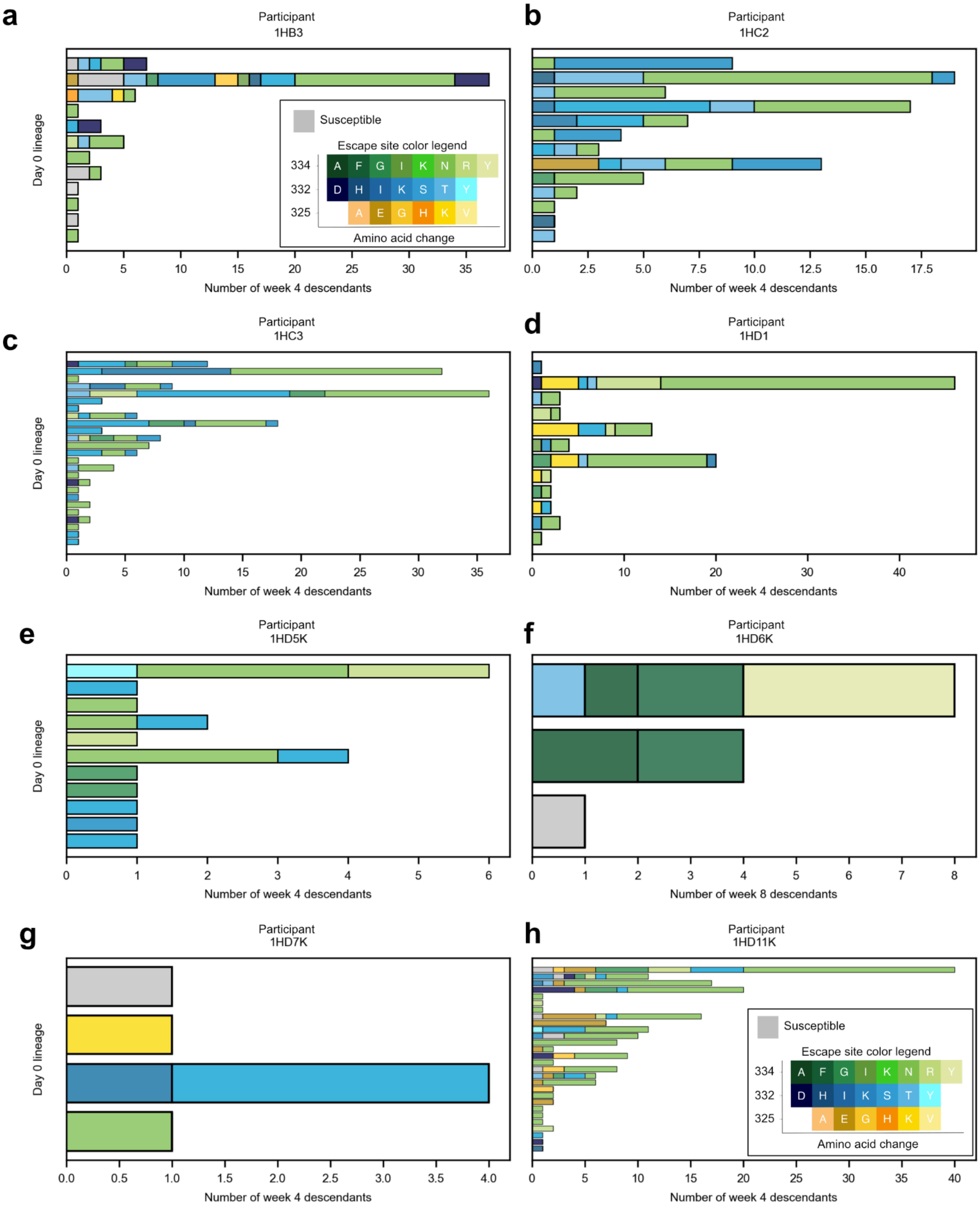

a
Participant
1HB3
Susceptible
Escape site color legend
334
A F G I K N R Y
332
D H I K S T Y
325
A E G H K V
Amino acid change
Day 0 lineage
Number of week 4 descendants
b
Participant
1HC2
Number of week 4 descendants
c
Participant
1HC3
Day 0 lineage
Number of week 4 descendants
d
Participant
1HD1
Number of week 4 descendants
e
Participant
1HD5K
Day 0 lineage
Number of week 4 descendants
f
Participant
1HD6K
Number of week 8 descendants
g
Participant
1HD7K
Day 0 lineage
Number of week 4 descendants
h
Participant
1HD11K
Susceptible
Escape site color legend
334
A F G I K N R Y
332
D H I K S T Y
325
A E G H K V
Amino acid change
Number of week 4 descendants

**Supplemental Figure 11.** All 10-1074 treatment responders show evidence of multiple resistant lineages coexisting during the trial. **(a-h)** Bar plots of the escape mutations carried by the week 4 descendants of each day 0 lineage. For participant 1HD6K, week 8 is plotted due to sample availability. Day 0 lineages are defined as the connected components of the participant's corresponding network plot (Supp. Fig. 10). Data for participants 1HD4K & 1HD10K are shown in figure 3.

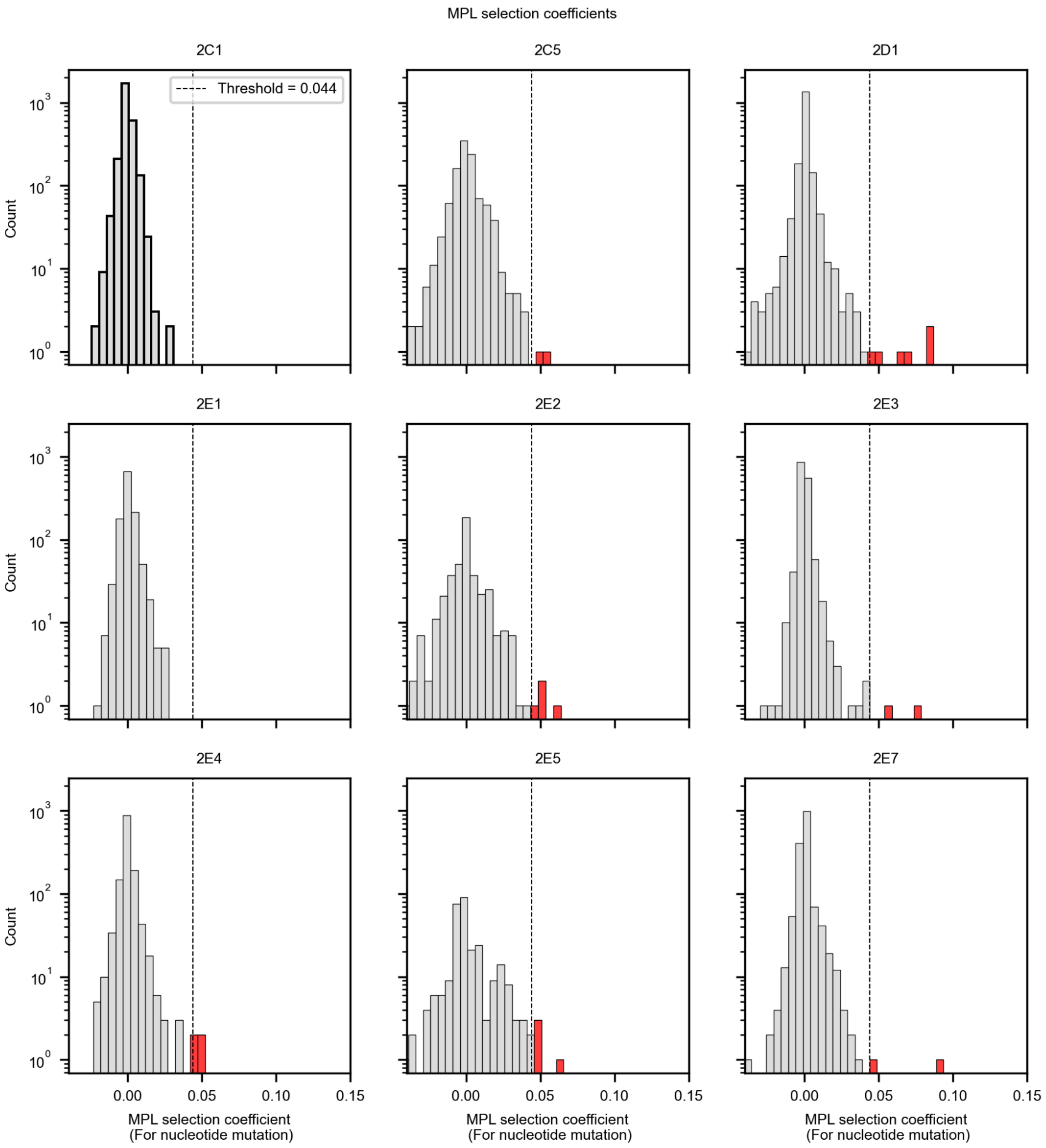


**Supplemental Figure 12.** Histograms of MPL selection coefficients inferred for all 3BNC117-treated study participants (Sohail et al., 2021). Sites with selection coefficients $s \geq 0.044$ were identified as strongly selected.

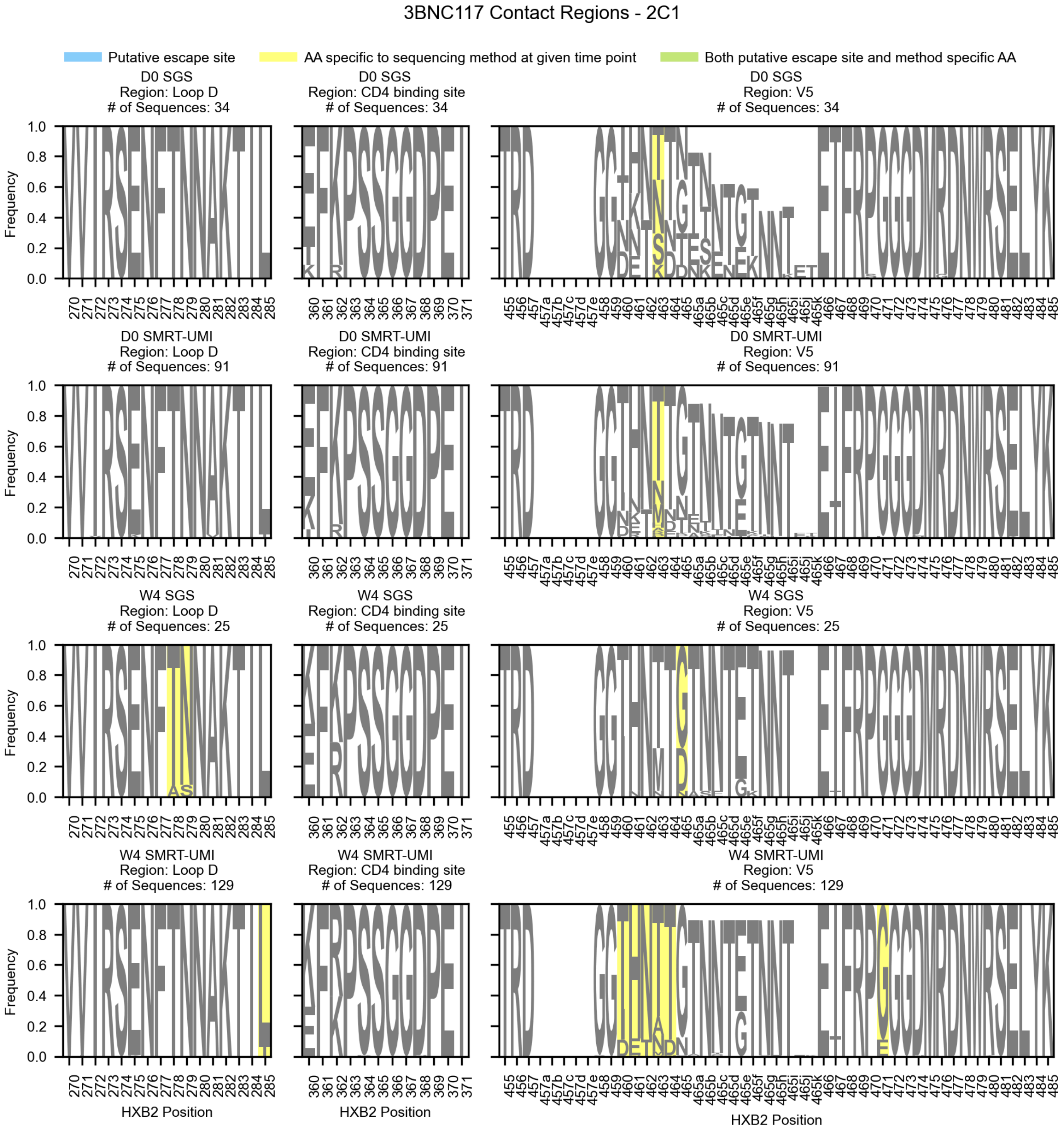


**Supplemental Figure 13.** Logo plots of amino acid frequencies in 3BNC117 contact regions for participant 2C1 during the trial. Rows show data for a given trial time point, day 0 (D0) or week 4 (W4), and sequencing method, single genome sequencing (SGS) from (Caskey et al., 2015; Schoofs et al., 2016) or SMRT-UMI sequencing collected here using the protocol from (Westfall et al., 2024). Columns represent distinct 3BNC117 contact regions (Schoofs et al., 2016; Zhou et al., 2013). Loci at which an AA was identified only by a single sequencing method are highlighted in yellow. Loci identified by one or more of our selection scans are highlighted in blue. Loci with both of these properties are highlighted in green.

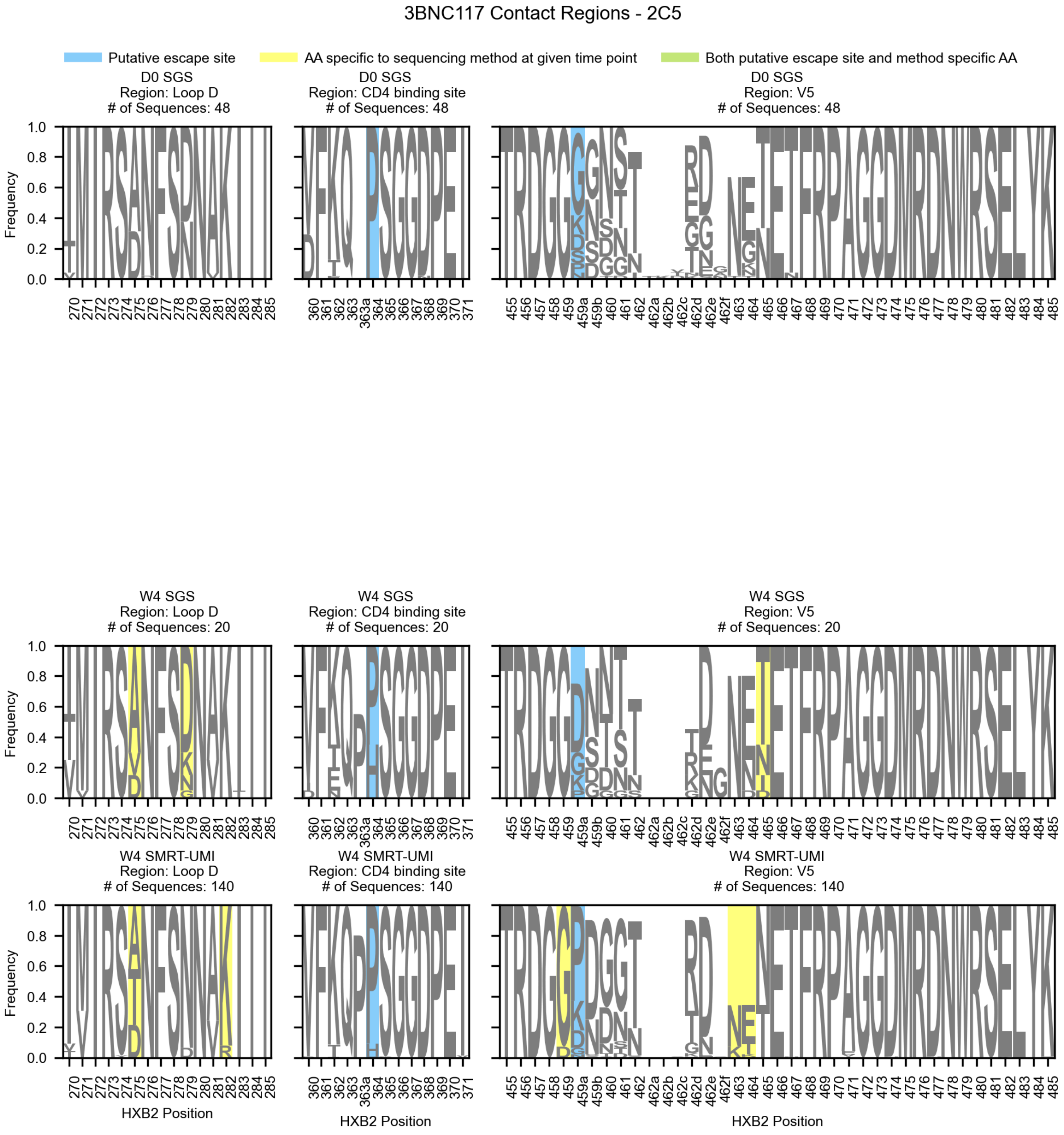


**Supplemental Figure 14.** Logo plots of amino acid frequencies in 3BNC117 contact regions for participant 2C5 during the trial. Rows show data for a given trial time point, day 0 (D0) or week 4 (W4), and sequencing method, single genome sequencing (SGS) from (Caskey et al., 2015; Schoofs et al., 2016) or SMRT-UMI sequencing collected here using the protocol from (Westfall et al., 2024). Columns represent distinct 3BNC117 contact regions (Schoofs et al., 2016; Zhou et al., 2013). Loci at which an AA was identified only by a single sequencing method are highlighted in yellow. Loci identified by one or more of our selection scans are highlighted in blue. Loci with both of these properties are highlighted in green. 2C5 had no day 0 SMRT-UMI sequencing resulting in a blank second row.

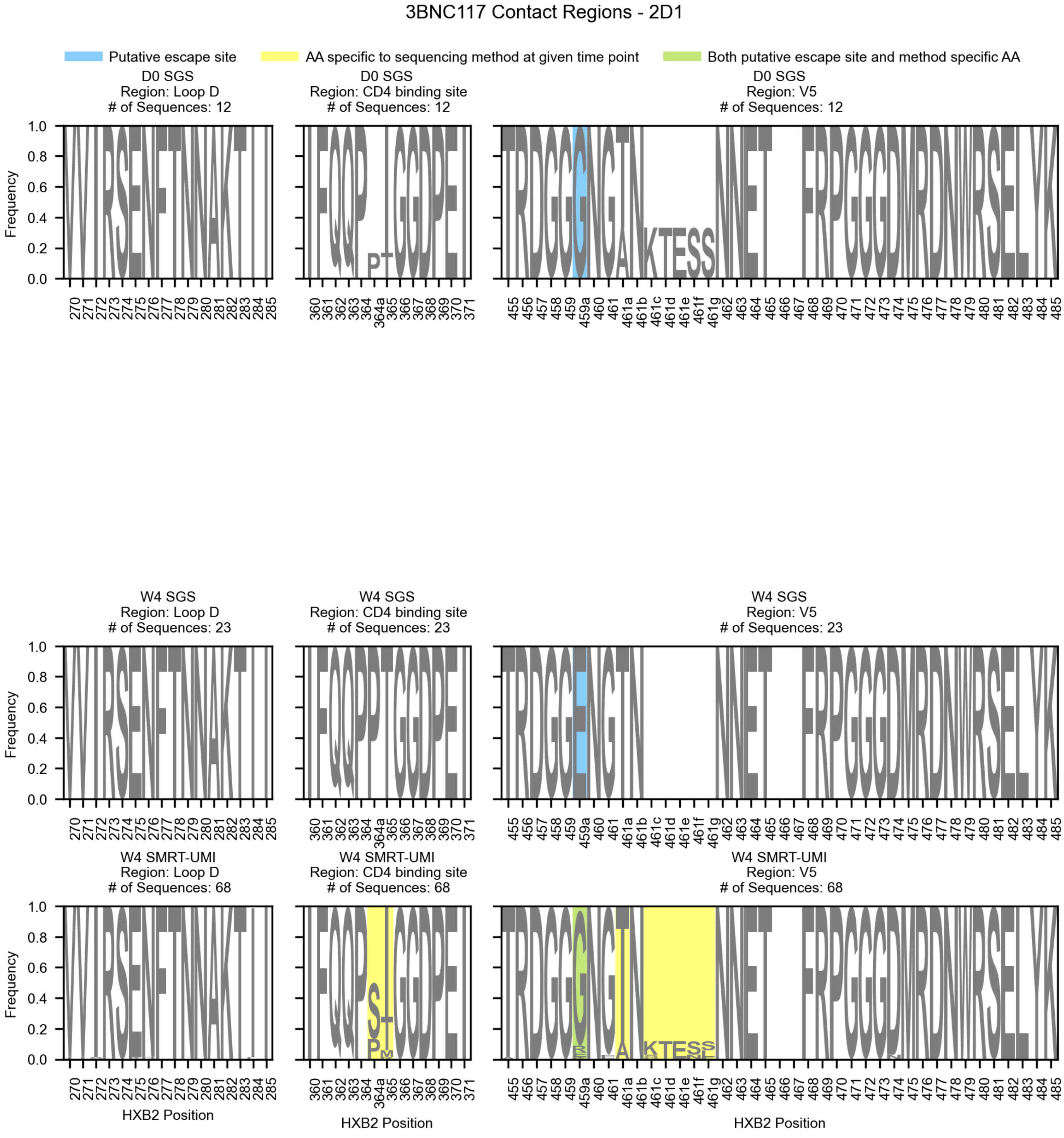


**Supplemental Figure 15.** Logo plots of amino acid frequencies in 3BNC117 contact regions for participant 2D1 during the trial. Rows show data for a given trial time point, day 0 (D0) or week 4 (W4), and sequencing method, single genome sequencing (SGS) from (Caskey et al., 2015; Schoofs et al., 2016) or SMRT-UMI sequencing collected here using the protocol from (Westfall et al., 2024). Columns represent distinct 3BNC117 contact regions (Schoofs et al., 2016; Zhou et al., 2013). Loci at which an AA was identified only by a single sequencing method are highlighted in yellow. Loci identified by one or more of our selection scans are highlighted in blue. Loci with both of these properties are highlighted in green. 2D1 had no day 0 SMRT-UMI sequencing resulting in a blank second row.

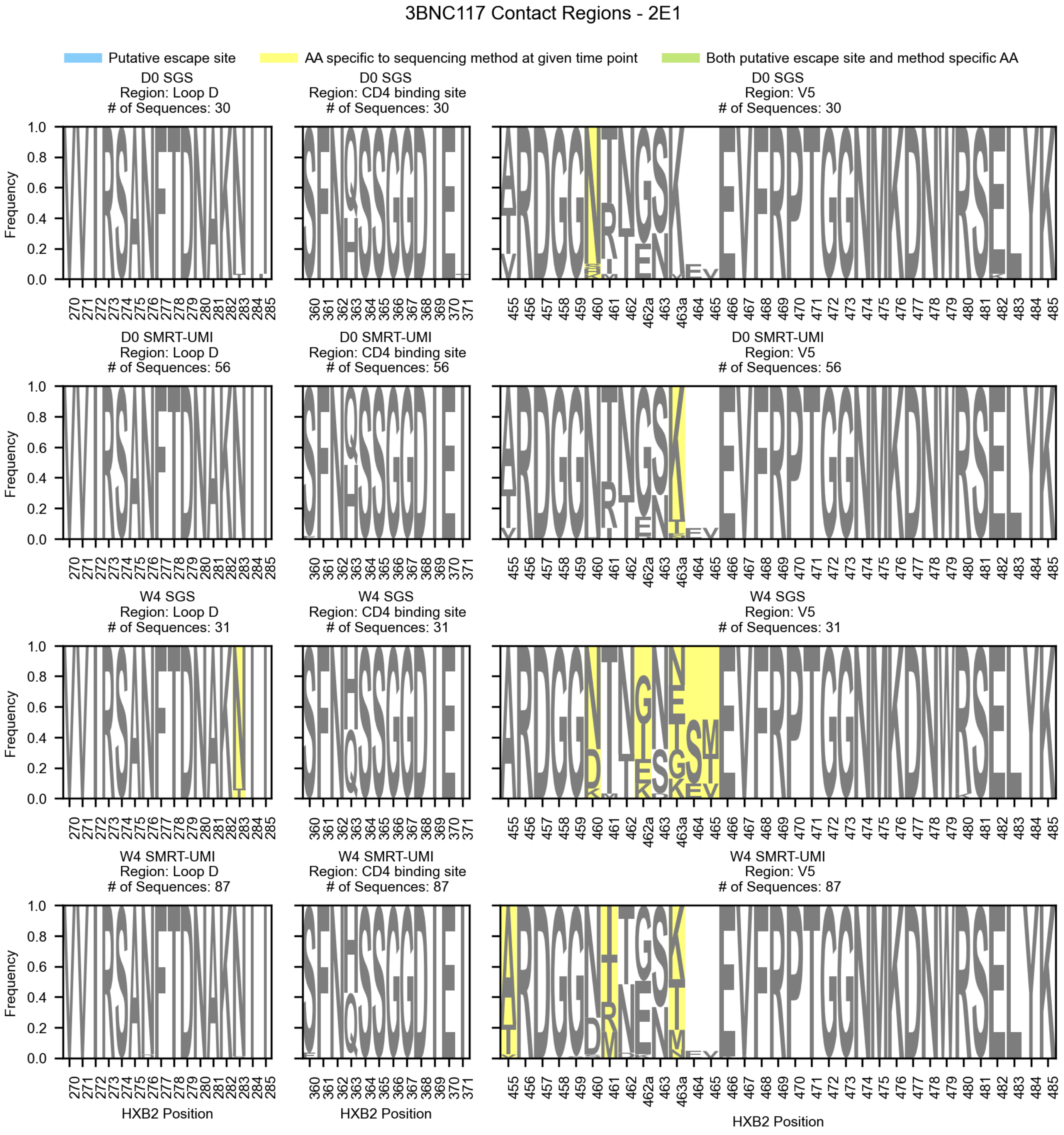


**Supplemental Figure 16.** Logo plots of amino acid frequencies in 3BNC117 contact regions for participant 2E1 during the trial. Rows show data for a given trial time point, day 0 (D0) or week 4 (W4), and sequencing method, single genome sequencing (SGS) from (Caskey et al., 2015; Schoofs et al., 2016) or SMRT-UMI sequencing collected here using the protocol from (Westfall et al., 2024). Columns represent distinct 3BNC117 contact regions (Schoofs et al., 2016; Zhou et al., 2013). Loci at which an AA was identified only by a single sequencing method are highlighted in yellow. Loci identified by one or more of our selection scans are highlighted in blue. Loci with both of these properties are highlighted in green.

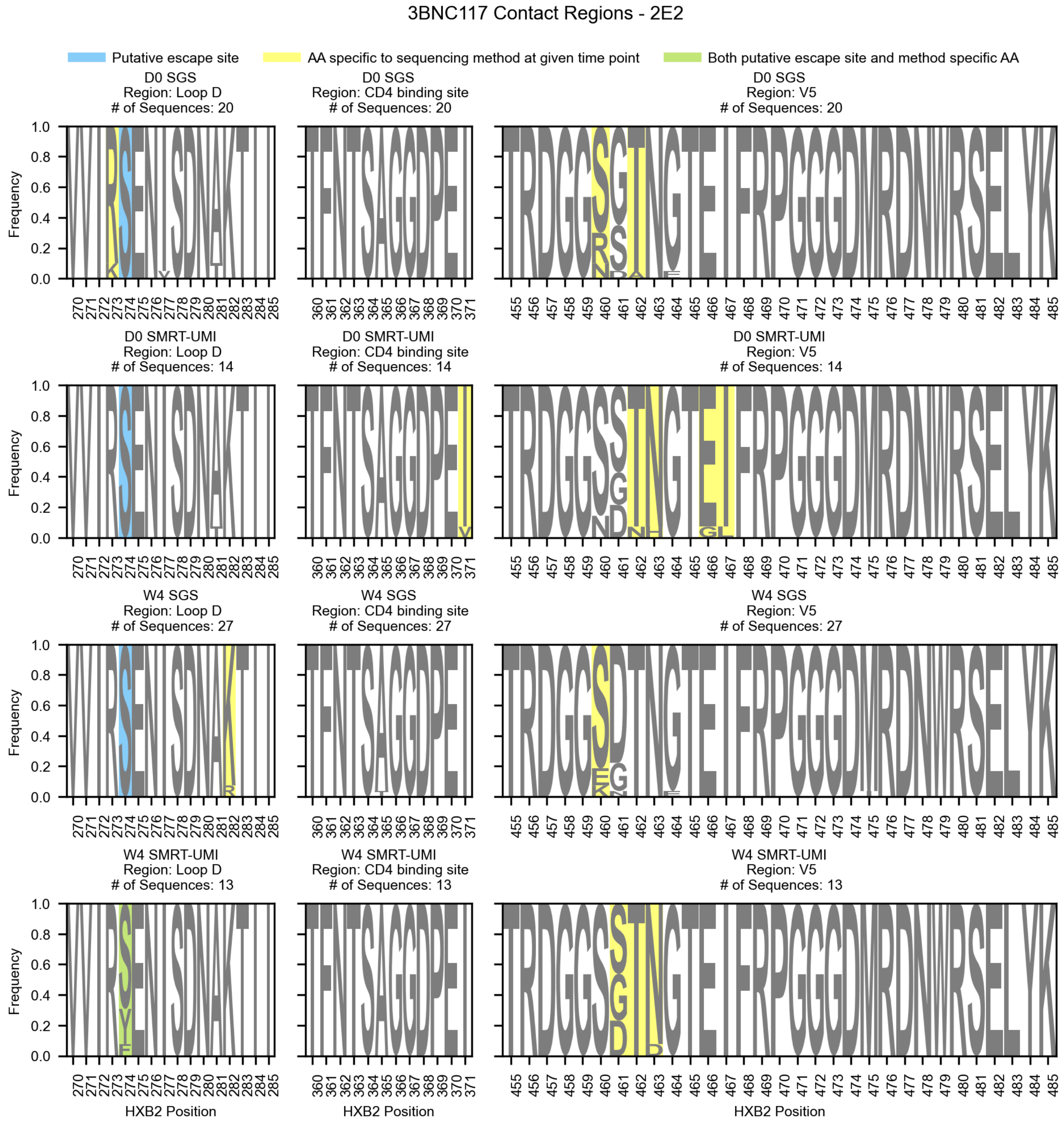


**Supplemental Figure 17.** Logo plots of amino acid frequencies in 3BNC117 contact regions for participant 2E2 during the trial. Rows show data for a given trial time point, day 0 (D0) or week 4 (W4), and sequencing method, single genome sequencing (SGS) from (Caskey et al., 2015; Schoofs et al., 2016) or SMRT-UMI sequencing collected here using the protocol from (Westfall et al., 2024). Columns represent distinct 3BNC117 contact regions (Schoofs et al., 2016; Zhou et al., 2013). Loci at which an AA was identified only by a single sequencing method are highlighted in yellow. Loci identified by one or more of our selection scans are highlighted in blue. Loci with both of these properties are highlighted in green.

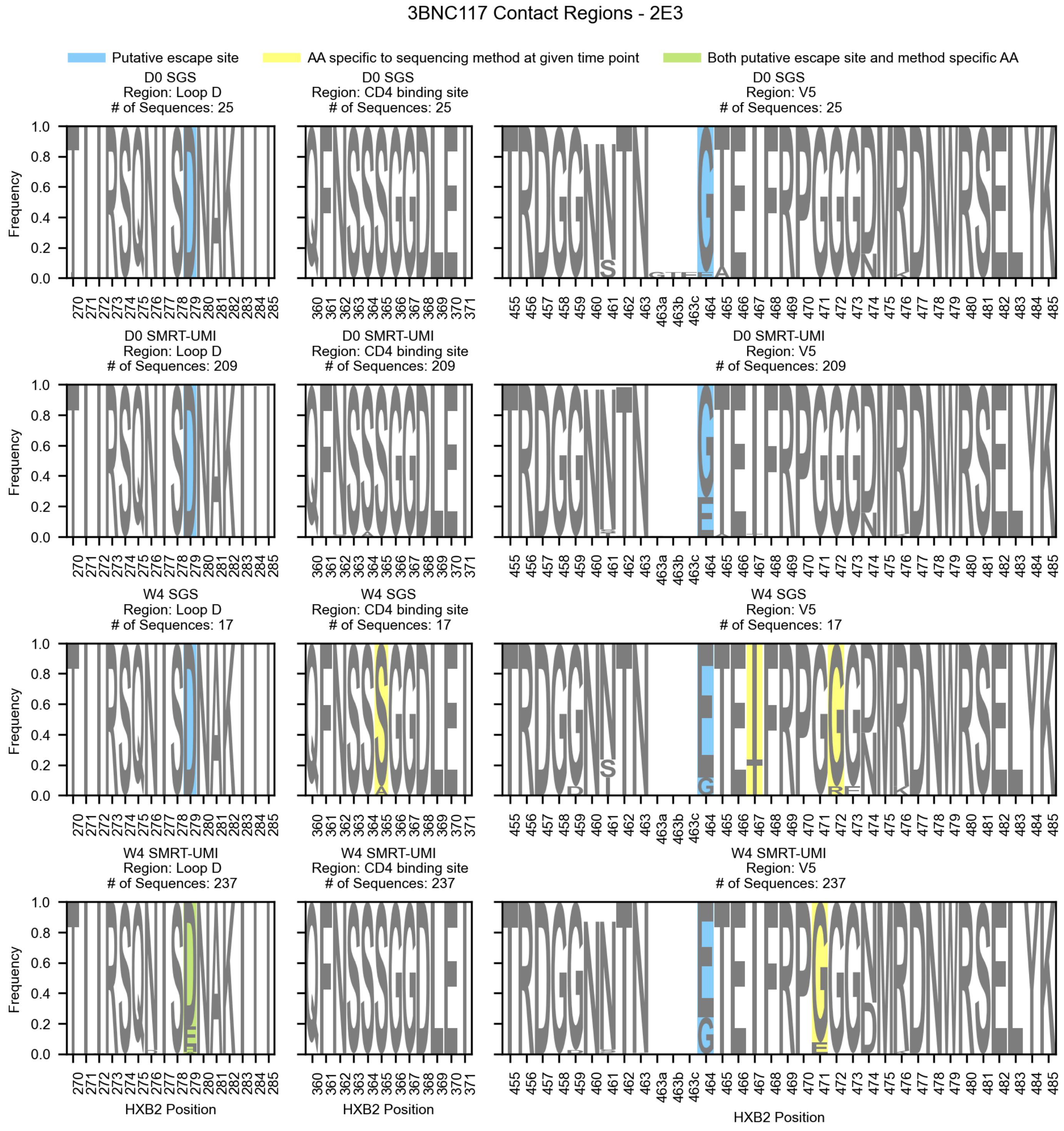


**Supplemental Figure 18.** Logo plots of amino acid frequencies in 3BNC117 contact regions for participant 2E3 during the trial. Rows show data for a given trial time point, day 0 (D0) or week 4 (W4), and sequencing method, single genome sequencing (SGS) from (Caskey et al., 2015; Schoofs et al., 2016) or SMRT-UMI sequencing collected here using the protocol from (Westfall et al., 2024). Columns represent distinct 3BNC117 contact regions (Schoofs et al., 2016; Zhou et al., 2013). Loci at which an AA was identified only by a single sequencing method are highlighted in yellow. Loci identified by one or more of our selection scans are highlighted in blue. Loci with both of these properties are highlighted in green.

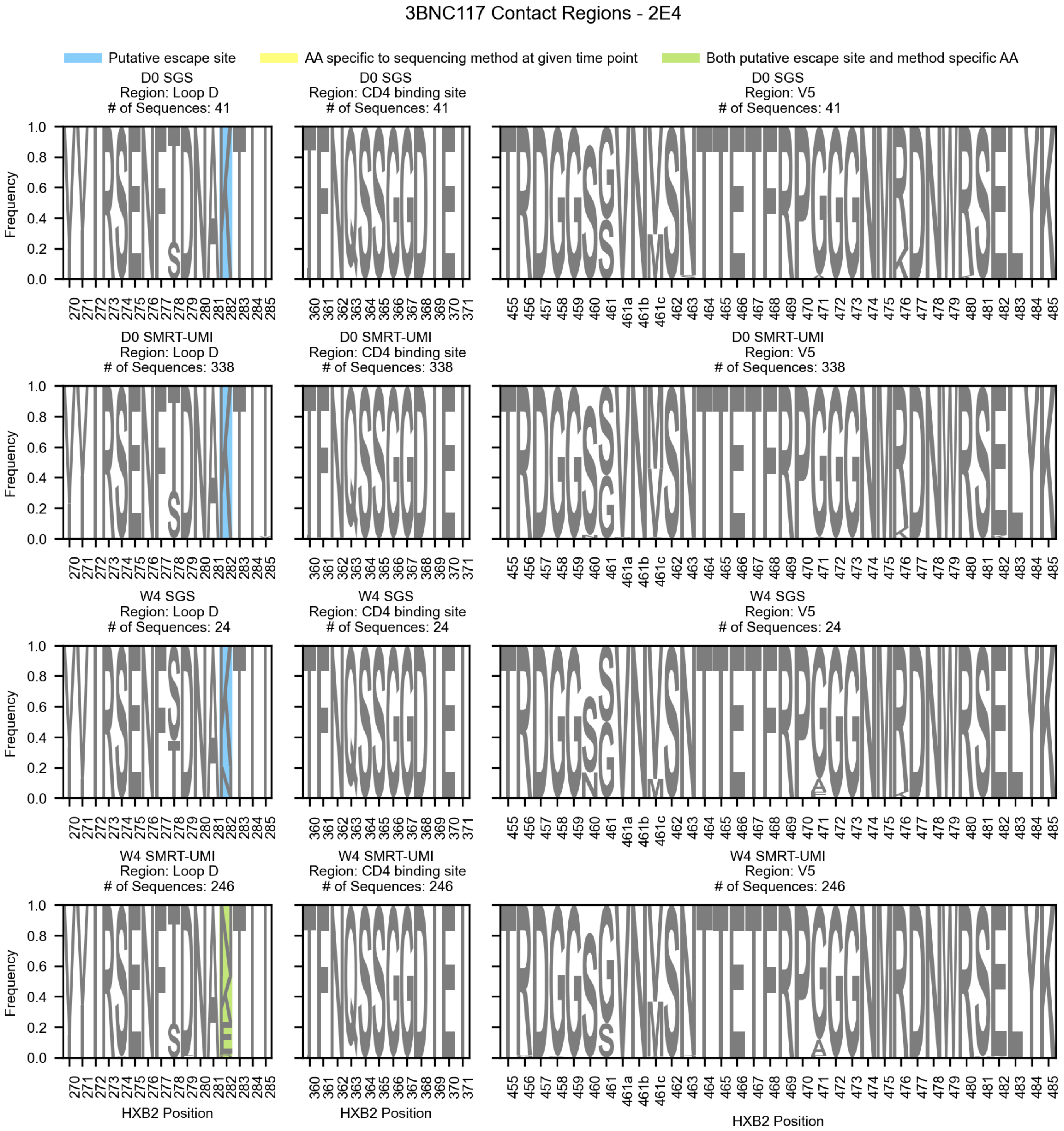


**Supplemental Figure 19.** Logo plots of amino acid frequencies in 3BNC117 contact regions for participant 2E4 during the trial. Rows show data for a given trial time point, day 0 (D0) or week 4 (W4), and sequencing method, single genome sequencing (SGS) from (Caskey et al., 2015; Schoofs et al., 2016) or SMRT-UMI sequencing collected here using the protocol from (Westfall et al., 2024). Columns represent distinct 3BNC117 contact regions (Schoofs et al., 2016; Zhou et al., 2013). Loci at which an AA was identified only by a single sequencing method are highlighted in yellow. Loci identified by one or more of our selection scans are highlighted in blue. Loci with both of these properties are highlighted in green.

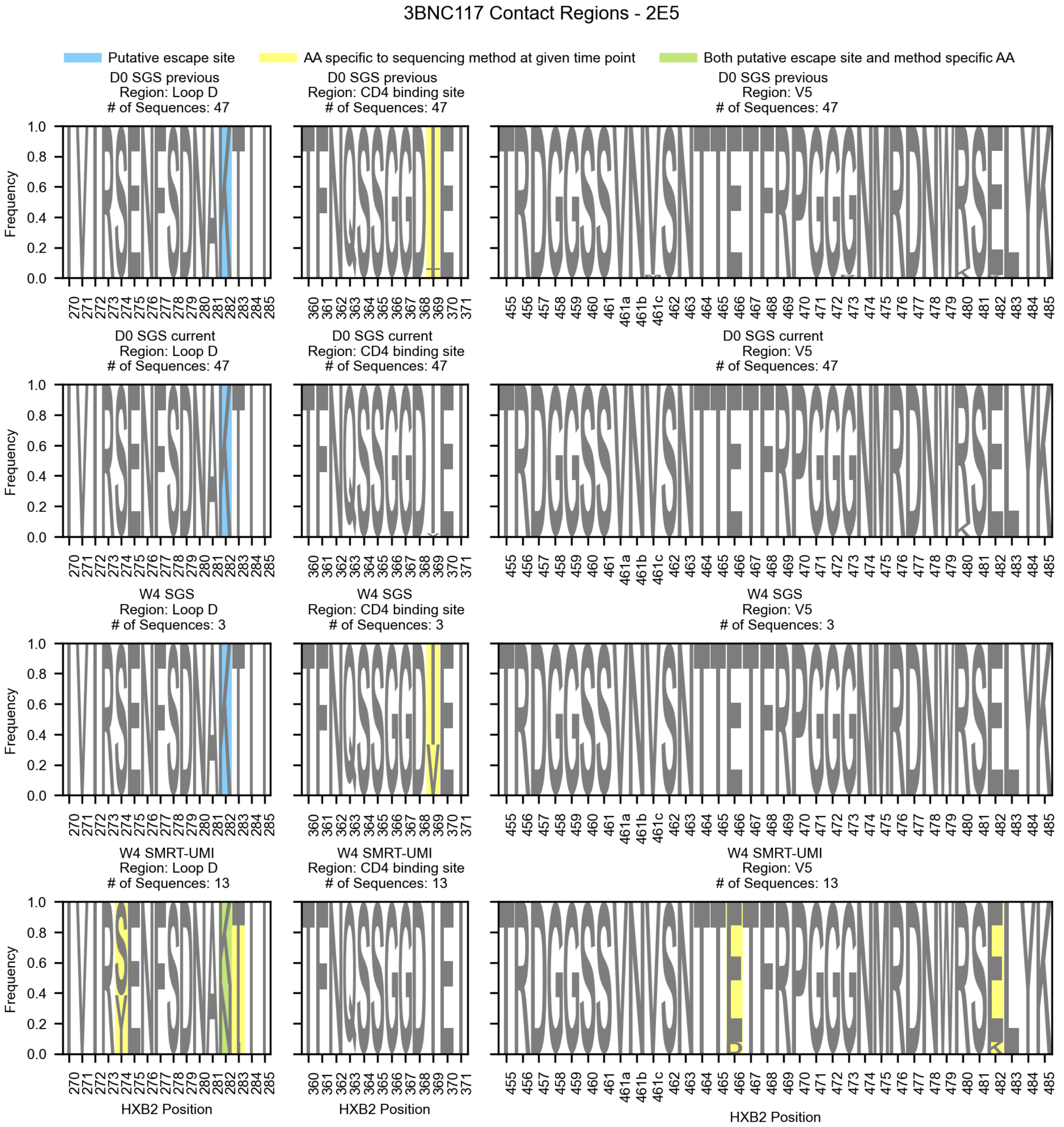


**Supplemental Figure 20.** Logo plots of amino acid frequencies in 3BNC117 contact regions for participant 2E5 during the trial. Rows show data for a given trial time point, day 0 (D0) or week 4 (W4), and sequencing method, single genome sequencing “SGS previous” sequenced by (Caskey et al., 2015; Schoofs et al., 2016), single genome sequencing “SGS current” sequenced here, or SMRT-UMI sequences collected here using the protocol from (Westfall et al., 2024). Columns represent distinct 3BNC117 contact regions (Schoofs et al., 2016; Zhou et al., 2013). Loci at which an AA was identified only by a single sequencing method are highlighted in yellow. Loci identified by one or more of  our selection scans are highlighted in blue. Loci with both of these properties are highlighted in green.

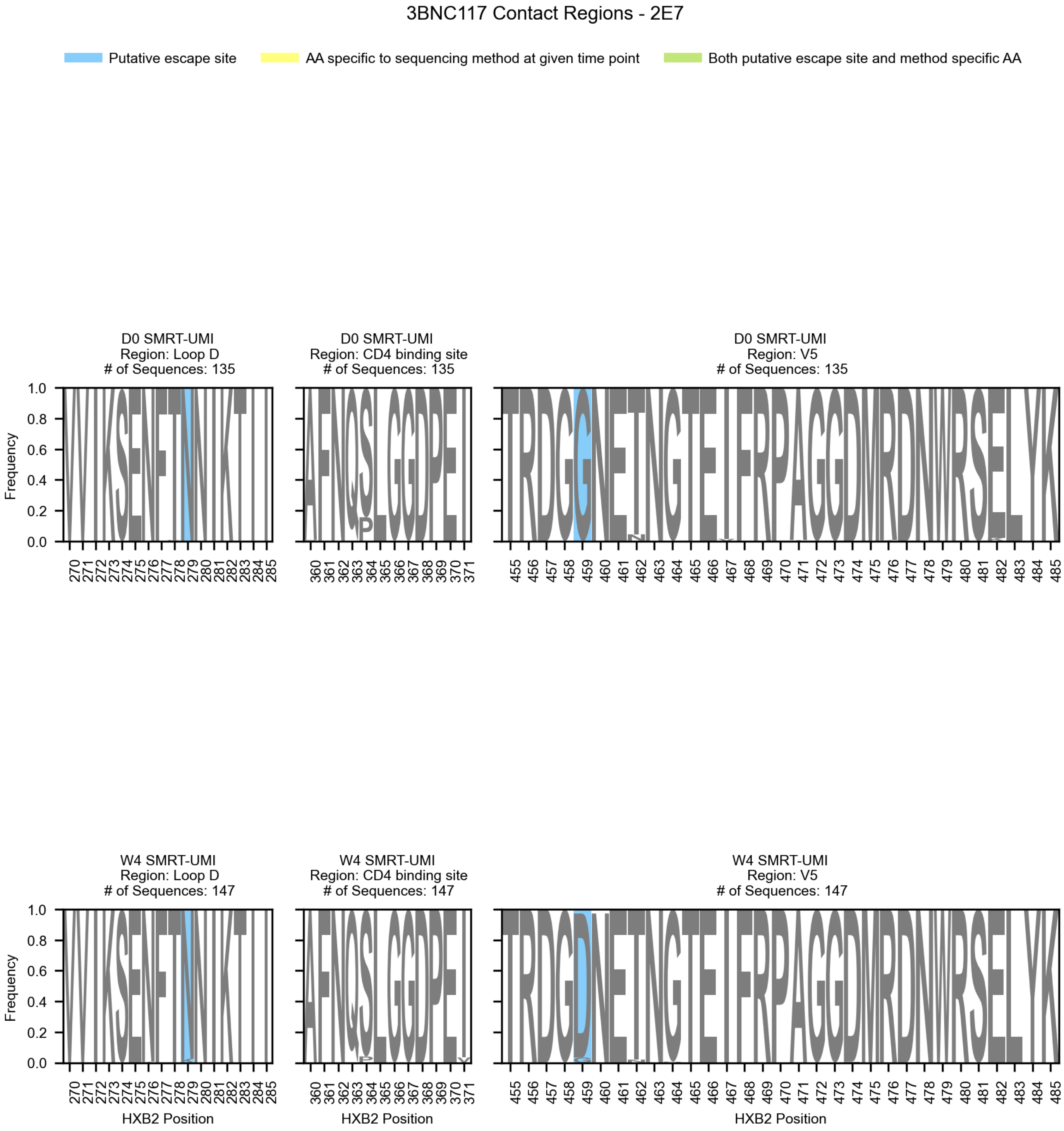


**Supplemental Figure 21.** Logo plots of amino acid frequencies in 3BNC117 contact regions for participant 2E7 during the trial. Rows show data for SMRT-UMI sequences collected here using the protocol from (Westfall et al., 2024) for a given trial time point, day 0 (D0) or week 4 (W4). Columns represent distinct 3BNC117 contact regions (Schoofs et al., 2016; Zhou et al., 2013). Loci identified by one or more of our selection scans are highlighted in blue. 2E7 had no SGS sequencing resulting in blank first and third rows.

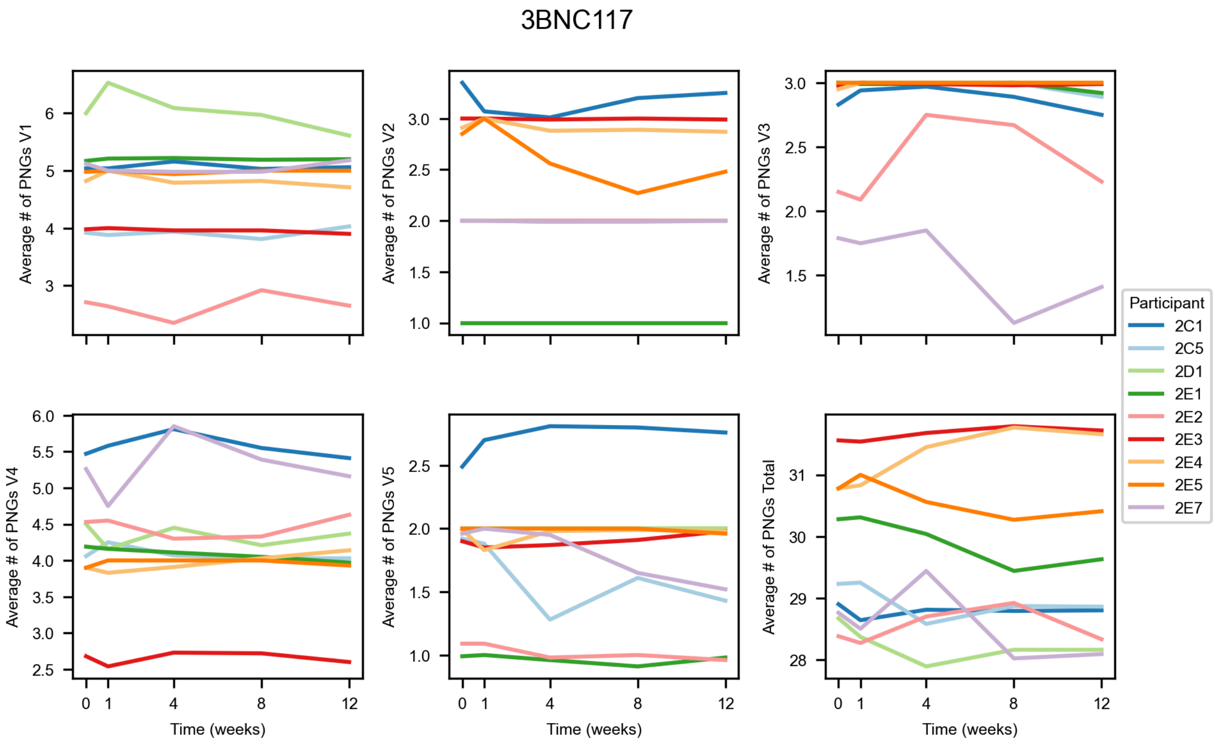


**Supplemental Figure 22.** Average numbers of potential N-linked glycosylation sites (PNGs) within each participant throughout the trial in the 3BNC117 treated cohort. Each panel displays data for a single variable loop (V1-V5) or the total count of PNGs summed across all variable loops.

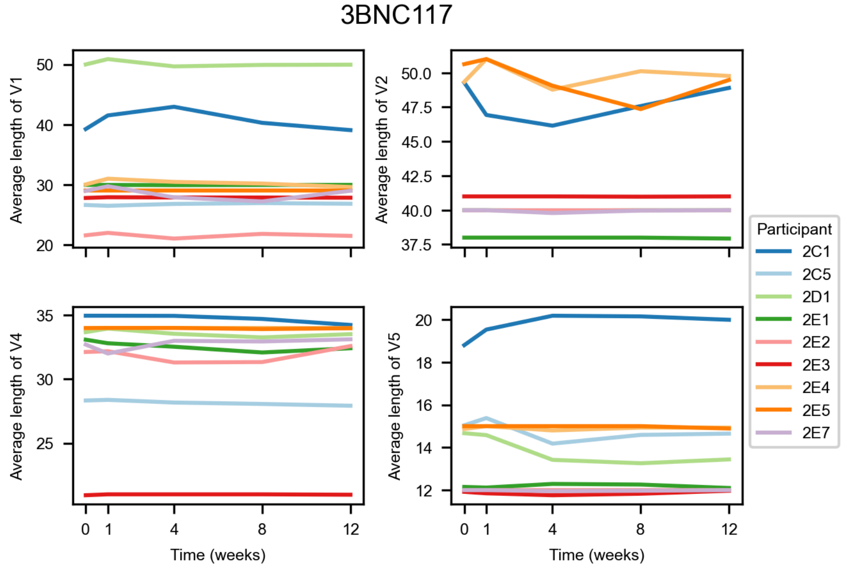


**Supplemental Figure 23.** Average lengths of variable loop regions in participants treated with 3BNC117. V3 is not shown as its length is highly conserved (De Wolf et al., 1994; Laakso et al., 2007; Zhong et al., 1995).

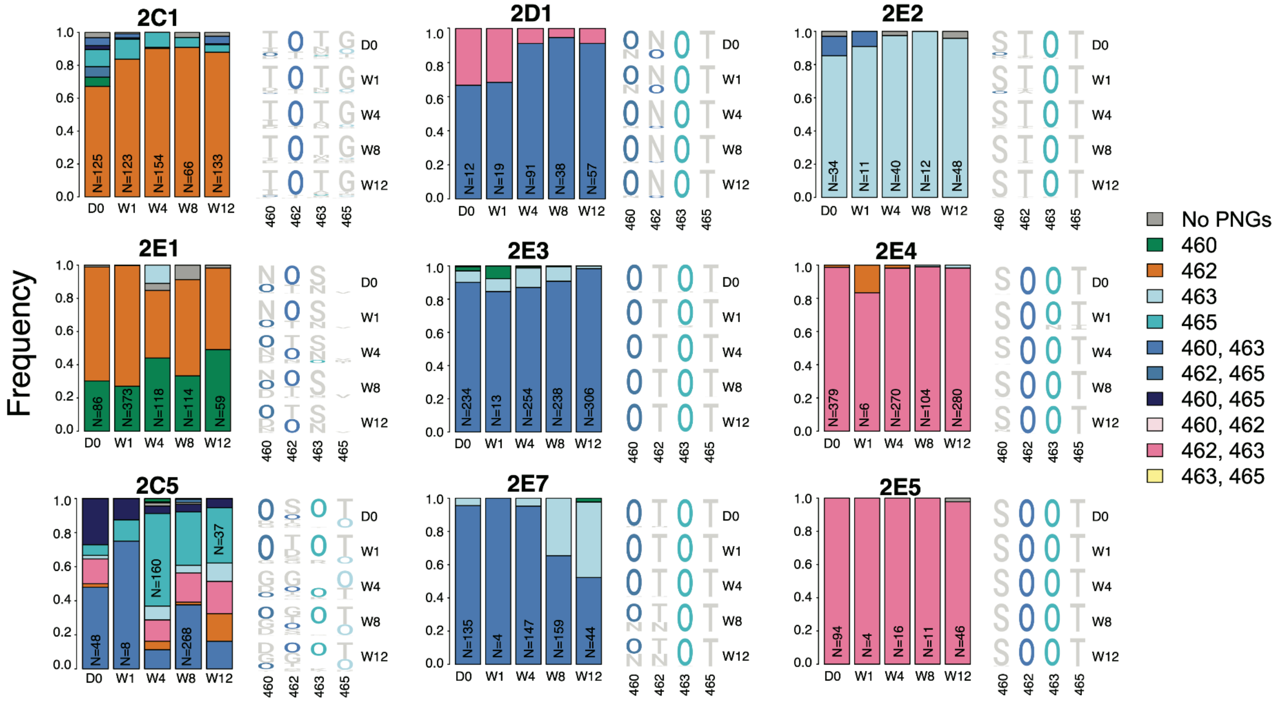


**Supplemental Figure 24.** Potential N-linked glycosylation (PNG) site patterns over time in the V5 region for participants treated with 3BNC117. **Left subpanel:** Stacked bar plots show the proportions of sequences carrying PNGs at loci in V5 at a given trial time point (D0, W1, W4, W8, or W12). Colors indicate which loci carry PNGs (listed in the legend). The total number of sequences available during the time point is labeled on each bar. **Right subpanel:** Logo plots of amino acid frequencies (with PNGs denoted by O) at V5 loci 460, 462, 463, and 465 during the trial.

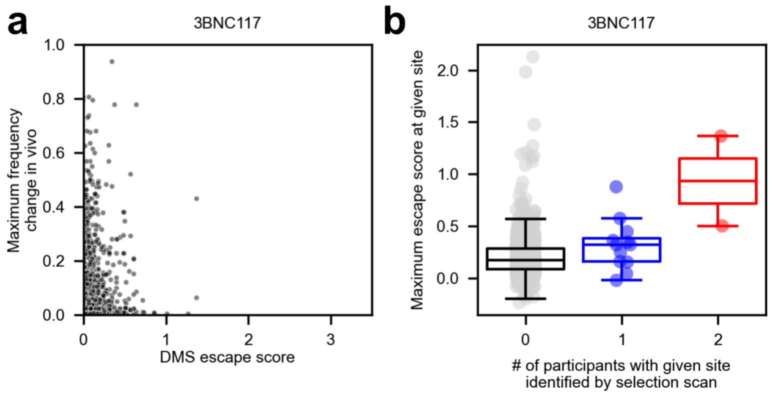


**Supplemental Figure 25.** Deep mutational scanning (DMS) escape scores against 3BNC117 for Env AAs on both TRO11 and BF520 backgrounds (Radford & Bloom, 2025) compared to their in vivo maximum frequency change in any 3BNC117-treated participant during the trial. Higher escape scores indicate less antibody neutralization by 3BNC117. Loci with no equivalent site in the DMS envelopes are omitted. **(b)** Box plots of maximum escape scores (across all AA identities and both DMS envelopes) for each locus stratified by the number of 3BNC117 participants in which the locus was identified as a putative escape site by one or more selection scan.

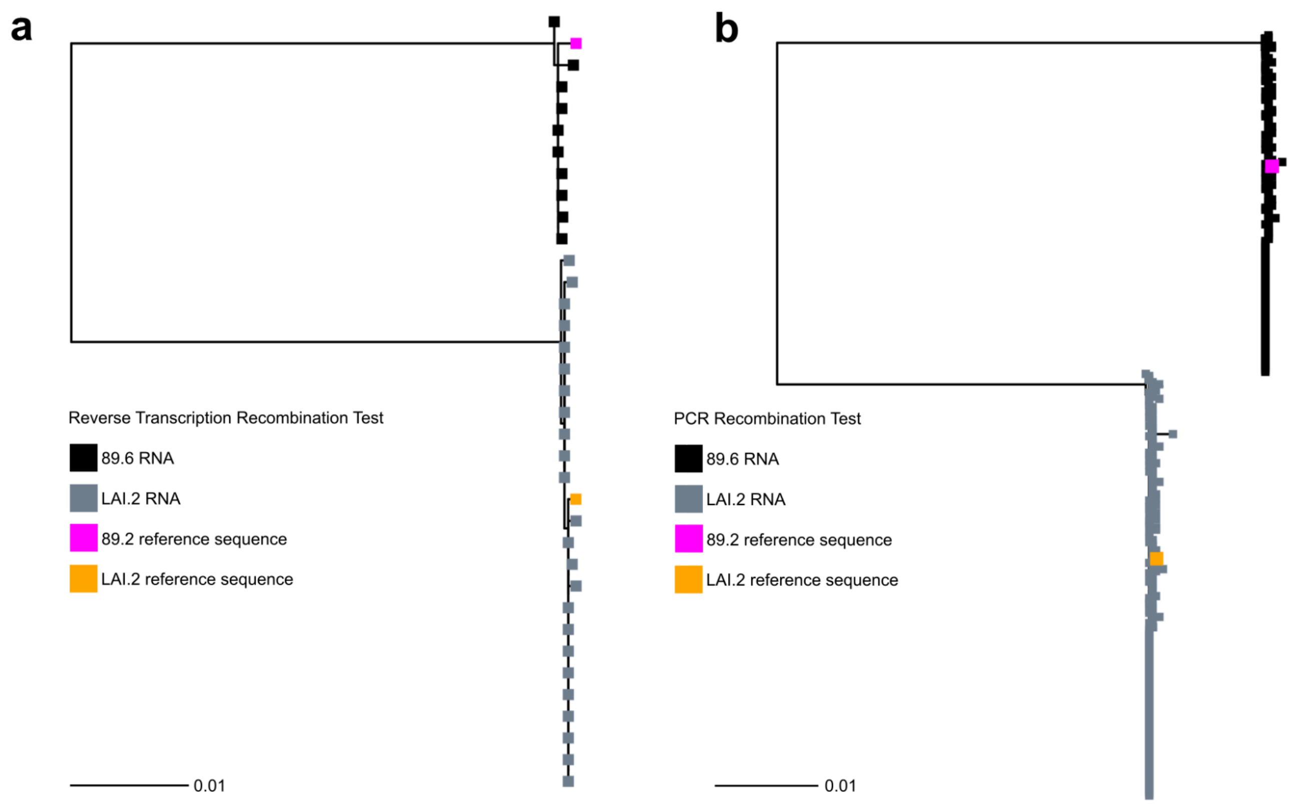


**Supplemental Figure 26:** Trees comparing RNA and cDNA sequences from separate molecular clones to analyze the occurrence of recombination in the SMRT-UMI protocol.

**(a)** RNA from molecular clones 89.6 (black) and LAI.2 (grey) was combined for cDNA synthesis and sequenced using single genome amplification. Compared to reference sequences, no cDNA recombinants were identified. Trees were made with IQTree2 software using a Tamura-Nei substitution model (Minh et al., 2020; Tamura & Nei, 1993) **(b)** 89.6 cDNA (black) and LAI.2 cDNA (grey) were labelled with different SMRT-UMI sample IDs,combined, and processed with bulk PCR amplification. PacBio long read sequencing was performed and no PCR recombinants were identified following PORPID pipeline filtering. Trees were made with IQTree2 software using a Tamura-Nei substitution model (Minh et al., 2020; Tamura & Nei, 1993).
