## Supplemental Tables for "Recurrent mutations drive rapid HIV escape from two broadly neutralizing antibodies *in vivo*"

#### Sampled 10-1074 timepoints

**Supplemental Table 1.**

| Participant | Timepoint sampled | Viral Load (c/mL) | Sequences Recovered using SMRT-UMI | Sequences analyzed from Caskey et al, 2017 | Total sequences analyzed |
|---|---|---|---|---|---|
| 1HB3 | Day 0 | 35810 | 41 | 29 | 70 |
| | Week 1 | 2990 | 12 | 16 | 28 |
| | Week 4 | 17380 | 40 | 28 | 68 |
| | Week 8 | 36580 | 116 | | 116 |
| 1HC2 | Day 0 | 77610 | 51 | 31 | 82 |
| | Week 1 | 1500 | 6 | 24 | 30 |
| | Week 4 | 49840 | 60 | 28 | 88 |
| 1HC3 | Day 0 | 20140 | 70 | 29 | 99 |
| | Week 1 | 520 | 18 | 30 | 48 |
| | Week 4 | 20800 | 128 | 32 | 160 |
| 1HD1 | Day 0 | 41770 | 140 | 32 | 172 |
| | Week 1 | 2900 | 23 | 19 | 42 |
| | Week 4 | 26710 | 76 | 24 | 100 |
| | Week 8 | 32940 | 198 | | 198 |
| 1HD10K | Day 0 | 10500 | 100 | 19 | 119 |
| | Week 1 | 496 | 3 | | 3 |
| | Week 4 | 503 | 21 | 12 | 33 |
| | Week 8 | 2670 | 28 | 20 | 48 |
| | Week 12 | 12900 | 87 | 23 | 110 |
| 1HD11K | Day 0 | 63400 | 115 | 36 | 151 |
| | Week 1 | 757 | 10 | | 10 |
| | Week 4 | 50500 | 159 | 31 | 190 |
| | Week 8 | 97000 | 105 | | 105 |
| | Week 12 | 224000 | 47 | | 47 |
| 1HD4K | Day 0 | 60300 | 136 | 23 | 159 |
| | Week 1 | 5140 | 10 | 20 | 30 |
| | Week 4 | 60000 | 52 | 33 | 85 |
| | Week 8 | 61700 | 110 | | 110 |
| | Week 12 | 47300 | 70 | | 70 |
| 1HD5K | Day 0 | 7570 | 27 | 23 | 50 |
| | Week 1 | 367 | | 5 | 5 |
| | Week 4 | 2960 | 1 | 19 | 20 |
| | Week 8 | 114000 | 146 | | 146 |

| | | | | | |
|---|---|---|---|---|---|
| 1HD6K | Day 0 | 10100 | 20 | 32 | 52 |
| | Week 8 | 992 | 4 | 9 | 13 |
| | Week 16 | 5900 | 11 | | 11 |
| 1HD7K | Day 0 | 9250 | 15 | | 15 |
| | Week 1 | 1170 | 15 | | 15 |
| | Week 4 | 5990 | 7 | | 7 |
| 1HD9K | Day 0 | 6960 | 14 | 30 | 44 |
| | Week 1 | 8120 | 12 | | 12 |
| | Week 4 | 18600 | 20 | | 20 |

Sampled 3BNC117 Timepoints

**Supplemental Table 2.**

| Participant | Timepoint Sampled | Viral Load (c/mL) | Sequences Recovered using SMRT-UMI | Sequences analyzed from Caskey et al, 2016 | Total sequences analyzed |
|---|---|---|---|---|---|
| 2C1 | Day 0 | 47650 | 91 | 34 | 125 |
| | Week 1 | 7470 | 123 | | 123 |
| | Week 4 | 25610 | 129 | 25 | 154 |
| | Week 8 | 32310 | 66 | | 66 |
| | Week 12 | 25150 | 133 | | 133 |
| 2C5 | Day 0 | 9260 | | 48 | 48 |
| | Week 1 | 410 | 8 | | 8 |
| | Week 4 | 8550 | 140 | 20 | 160 |
| | Week 8 | 12840 | 268 | | 268 |
| | Week 12 | 14130 | 15 | 22 | 37 |
| 2D1 | Day 0 | 53470 | | 12 | 12 |
| | Week 1 | 5980 | 19 | | 19 |
| | Week 4 | 10960 | 68 | 23 | 91 |
| | Week 8 | 9820 | 38 | | 38 |
| | Week 12 | 8710 | 57 | | 57 |
| 2E1 | Day 0 | 15780 | 56 | 30 | 86 |
| | Week 1 | 404 | 373 | | 373 |
| | Week 4 | 8159 | 87 | 31 | 118 |
| | Week 8 | 13486 | 114 | | 114 |
| | Week 12 | 7446 | 32 | 27 | 59 |
| 2E2 | Day 0 | 6990 | 14 | 20 | 34 |
| | Week 1 | 1663 | 11 | | 11 |
| | Week 4 | 4273 | 13 | 27 | 40 |
| | Week 8 | 3465 | 12 | | 12 |
| | Week 12 | 1282 | 23 | 25 | 48 |
| 2E3 | Day 0 | 22030 | 209 | 25 | 234 |
| | Week 1 | 1308 | 13 | | 13 |
| | Week 4 | 33370 | 237 | 17 | 254 |
| | Week 8 | 27998 | 238 | | 238 |
| | Week 12 | 27517 | 282 | 24 | 306 |
| 2E4 | Day 0 | 32220 | 338 | 41 | 379 |
| | Week 1 | 3642 | 6 | | 6 |
| | Week 4 | 18005 | 246 | 24 | 270 |
| | Week 8 | 22100 | 104 | | 104 |
| | Week 12 | 61200 | 255 | 25 | 280 |
| 2E5 | Day 0 | 3610 | 47 | 27 | 94 |
| | Week 1 | 181 | | 4 | 4 |

| | | | | | |
|---|---|---|---|---|---|
| | Week 4 | 148 | | 16 | 16 |
| | Week 8 | 589 | 11 | | 11 |
| | Week 12 | 4770 | 23 | 23 | 46 |
| 2E7 | Day 0 | 11730 | 135 | | 135 |
| | Week 1 | 690 | 4 | | 4 |
| | Week 4 | 67700 | 147 | | 147 |
| | Week 8 | 36800 | 159 | | 159 |

### 10-1074 accession numbers from previously reported sequences

**Supplemental Table 3.**
**10-1074 Single Genome Amplification Sequences Used (Caskey et al., 2017)**

| *Participant* | *NCBI Accession Numbers* |
|---|---|
| *1HB3* | KY323883-KY323911; KY323934-KYKY32349; KY3239690-KY323996 |
| *1HC2* | KY324081-KY324165 |
| *1HC3* | KY324166-KY324256 |
| *1HD1* | KY324398-KY324448; KY324479-KY324502 |
| *1HD4K* | KY324517-KY324592 |
| *1HD5K* | KY324593-KY324639 |
| *1HD6K* | KY324640-KY324671; KY324726-KY324734 |
| *1HD7K* | No prev. sequencing |
| *1HD9K* | KY324805-KY324834 |
| *1HD10K* | KY324257-KY324330 |
| *1HD11K* | KY324331-KY324397 |

### 3BNC117 accession numbers from previously reported sequences

**Supplemental Table 4.**
**3BNC117 Single Genome Amplification Sequences Used (Caskey et al., 2015; Schoofs et al., 2016)**

| *Participant* | *NCBI Accession Numbers* |
|---:|---|
| *2C1* | KX016905- KX016963 |
| *2C5* | KX028066-KX028135; KX028168-KX028187 |
| *2D1* | KX017008- KX17042 |
| *2E1* | KX028188-KX028244; KX28282-KX28312 |
| *2E2* | KX028313-KX28357; KX028385-KX028411 |
| *2E3* | KX028412-KX028460; KX028482-KX028498 |
| *2E4* | KX028499-KX028565; KX028595-KX028619 |
| *2E5* | KX028620-KX028689; KX028712-KX028714 |
| *2E7* | No prev. sequencing |

#### 10-1074 Variable Sites

**Supplemental Table 5.**
Variable loci within 10-1074 treated participants as assessed by 40% loss of the day 0 majority amino acid (see Materials & Methods). Loci which varied in three or more participants are included. Corresponding % losses of the majority AA are shown for each participant with variation in the corresponding “Max loss (%) by participant” list. Maximum DMS escape score indicates the maximum escape score of all amino acids assayed at the given site by Radford & Bloom 2025.

| HXB2 Coordinate | Participant count | Participants with variation | Max loss (%) by participant | Variable Loop | Maximum DMS escape score (Radford & Bloom 2025) |
|---|---|---|---|---|---|
| 334 | 9 | ['1HB3', '1HC2', '1HC3', '1HD1', '1HD4K', '1HD5K', '1HD6K', '1HD7K', '1HD11K'] | ['49.7%', '45.5%', '58.3%', '73.0%', '69.8%', '64.3%', '84.6%', '46.7%', '77.1%'] | FALSE | 3.536 |
| 332 | 5 | ['1HC2', '1HC3', '1HD4K', '1HD5K', '1HD7K'] | ['51.1%', '46.2%', '82.4%', '42.0%', '54.1%'] | FALSE | 3.586 |
| 406 | 4 | ['1HB3', '1HD4K', '1HD7K', '1HD10K'] | ['41.9%', '82.3%', '77.8%', '51.3%'] | TRUE | 0.4708 |
| 347 | 3 | ['1HD6K', '1HD7K', '1HD10K'] | ['100.0%', '54.1%', '46.8%'] | FALSE | 0.4044 |
| 397 | 3 | ['1HD4K', '1HD7K', '1HD10K'] | ['82.0%', '41.6%', '60.6%'] | TRUE | 0.2856 |
| 400 | 3 | ['1HD7K', '1HD10K', '1HD11K'] | ['87.5%', '61.8%', '71.5%'] | TRUE | 0.3084 |
| 409 | 3 | ['1HD6K', '1HD7K', '1HD10K'] | ['44.2%', '87.5%', '60.6%'] | TRUE | 0.4386 |
| 410 | 3 | ['1HD4K', '1HD7K', '1HD11K'] | ['67.9%', '52.4%', '62.8%'] | TRUE | 0.1435 |
| 464 | 3 | ['1HD4K', '1HD7K', '1HD10K'] | ['44.6%', '67.0%', '44.0%'] | TRUE | 0.2465 |
| 620 | 3 | ['1HD6K', '1HD7K', '1HD10K'] | ['100.0%', '80.0%', '50.6%'] | FALSE | 0.1883 |

#### 10-1074 MPL Identified sites

**Supplemental Table 6.**
All loci identified by MPL (Sohail et al., 2021) analysis in the 10-1074 treated cohort (see Materials & Methods). Column “s_MPL” indicates the selection coefficient inferred by MPL. Maximum DMS escape score indicates the maximum escape score of all amino acids assayed at the given site by Radford & Bloom 2025.

| participant | HXB2 Coordinate | s_MPL | Maximum DMS escape score (Radford & Bloom 2025) |
|---|---|---|---|
| **1HB3** | 332 | 0.044 | 3.586 |
| **1HB3** | 334 | 0.047 | 3.536 |
| **1HC2** | 332 | 0.064 | 3.586 |
| **1HC2** | 334 | 0.088 | 3.536 |
| **1HC3** | 332 | 0.062 | 3.586 |
| **1HC3** | 334 | 0.092 | 3.536 |
| **1HD1** | 325 | 0.045 | 2.562 |
| **1HD1** | 332 | 0.045 | 3.586 |
| **1HD1** | 334 | 0.069 | 3.536 |
| **1HD4K** | 328 | 0.059 | 1.705 |
| **1HD4K** | 334 | 0.09 | 3.536 |
| **1HD5K** | 332 | 0.044 | 3.586 |
| **1HD5K** | 334 | 0.047 | 3.536 |
| **1HD11K** | 334 | 0.066 | 3.536 |

### 10-1074 Escape Origin Quantification

**Supplemental Table 7.**
Quantification of lineage analysis results presented in Figure 3h,i and Supplemental Figure 11. Here we sum and report the number of unique combinations of escape AA and lineage (unique escape origins) we observed compared to the number of unique escape AAs without accounting for lineage (See Methods).

| Participant | # of unique escape AA and lineage combinations | # of unique escape AAs without accounting for lineage | Factor increase | # of additional origins |
|---|---|---|---|---|
| **1HB3** | 28 | 13 | 2.15 | 15 |
| **1HC2** | 32 | 8 | 4 | 24 |
| **1HC3** | 54 | 8 | 6.75 | 46 |
| **1HD1** | 32 | 10 | 3.2 | 22 |
| **1HD5K** | 15 | 6 | 2.5 | 9 |
| **1HD6K** | 6 | 4 | 1.5 | 2 |
| **1HD7K** | 4 | 4 | 1 | 0 |
| **1HD11K** | 60 | 12 | 5 | 48 |
| **1HD4K** | 15 | 6 | 2.5 | 9 |
| **1HD10K** | 15 | 6 | 2.5 | 9 |

### 3BNC117 Variable Sites

**Supplemental Table 8.**
Variable loci within 3BNC117 treated participants as assessed by >5% loss of the day 0 majority amino acid (see Materials & Methods). Loci which varied in three or more participants are included. Corresponding % losses of the majority AA are shown for each participant with variation in the corresponding “Max loss (%) by participant” list.

| HXB2 Coordinate | Participant count | Participants with variation | Max loss (%) by participant | Variable Loop |
|---|---|---|---|---|
| 282 | 4 | ['2E4', '2E5', '2C5', '2E2'] | ['91.3%', '37.5%', '6.9%', '8.3%'] | FALSE |
| 413 | 4 | ['2E7', '2E2', '2D1', '2E1'] | ['22.4%', '17.4%', '55.8%', '23.7%'] | TRUE |
| 404 | 4 | ['2E7', '2E4', '2C1', '2D1'] | ['7.9%', '17.2%', '9.0%', '5.3%'] | TRUE |
| 405 | 4 | ['2E7', '2E4', '2E2', '2C1'] | ['37.5%', '16.6%', '55.7%', '65.2%'] | TRUE |
| 145 | 4 | ['2E7', '2E5', '2D1', '2E3'] | ['73.1%', '6.2%', '21.1%', '31.5%'] | TRUE |
| 344 | 4 | ['2C5', '2E2', '2C1', '2D1'] | ['77.3%', '65.5%', '62.9%', '5.3%'] | FALSE |
| 462 | 4 | ['2E7', '2C5', '2E2', '2E1'] | ['30.2%', '23.3%', '7.3%', '25.9%'] | TRUE |
| 461 | 4 | ['2E4', '2D1', '2E1', '2E3'] | ['88.1%', '13.2%', '31.5%', '7.4%'] | TRUE |
| 337 | 4 | ['2E4', '2C5', '2C1', '2D1'] | ['54.6%', '22.6%', '17.4%', '25.0%'] | FALSE |
| 460 | 4 | ['2E4', '2C5', '2E1', '2E3'] | ['7.6%', '81.4%', '28.2%', '7.9%'] | TRUE |
| 410 | 3 | ['2E2', '2D1', '2E1'] | ['41.8%', '78.9%', '86.3%'] | TRUE |
| 29 | 3 | ['2E4', '2E5', '2D1'] | ['13.5%', '31.2%', '21.1%'] | FALSE |
| 400 | 3 | ['2C5', '2D1', '2E1'] | ['76.0%', '78.9%', '19.8%'] | TRUE |
| 463 | 3 | ['2C5', '2E1', '2E3'] | ['47.7%', '19.9%', '7.3%'] | TRUE |
| 471 | 3 | ['2E4', '2C1', '2E3'] | ['10.7%', '8.4%', '7.1%'] | FALSE |
| 829 | 3 | ['2E5', '2C5', '2E1'] | ['6.2%', '19.3%', '6.7%'] | FALSE |
| 832 | 3 | ['2E7', '2C5', '2E3'] | ['7.6%', '47.8%', '6.6%'] | FALSE |
| 403 | 3 | ['2E7', '2D1', '2E1'] | ['68.6%', '21.1%', '46.8%'] | TRUE |
| 396 | 3 | ['2C5', '2D1', '2E1'] | ['25.0%', '5.3%', '11.4%'] | TRUE |
| 398 | 3 | ['2C1', '2D1', '2E1'] | ['8.5%', '13.2%', '24.6%'] | TRUE |
| 32 | 3 | ['2E4', '2E1', '2E3'] | ['70.1%', '9.5%', '28.2%'] | FALSE |
| 347 | 3 | ['2E7', '2C5', '2C1'] | ['28.2%', '74.1%', '8.5%'] | FALSE |

| | | | | |
|---|---|---|---|---|
| 339 | 3 | ['2E7', '2C5', '2C1'] | ['8.2%', '34.8%', '65.4%'] | FALSE |
| 336 | 3 | ['2C5', '2C1', '2D1'] | ['25.0%', '63.9%', '49.5%'] | FALSE |
| 289 | 3 | ['2E4', '2E5', '2C1'] | ['22.3%', '6.2%', '12.1%'] | FALSE |
| 279 | 3 | ['2E7', '2C5', '2E3'] | ['33.5%', '93.4%', '18.9%'] | FALSE |
| 268 | 3 | ['2E5', '2C5', '2C1'] | ['6.2%', '18.6%', '21.4%'] | FALSE |
| 236 | 3 | ['2E5', '2D1', '2E1'] | ['6.2%', '5.3%', '11.4%'] | FALSE |
| 146 | 3 | ['2E7', '2C5', '2D1'] | ['73.1%', '26.8%', '5.3%'] | TRUE |
| 134 | 3 | ['2E7', '2C5', '2E2'] | ['31.1%', '19.0%', '42.3%'] | TRUE |
| 87 | 3 | ['2E7', '2C1', '2E3'] | ['64.5%', '37.9%', '11.2%'] | FALSE |
| 836 | 3 | ['2E7', '2C5', '2E2'] | ['64.1%', '23.1%', '6.3%'] | FALSE |

### 3BNC117 MPL & Multiple encoding analysis sites

**Supplemental Table 9.**
All loci identified by MPL (Sohail et al., 2021) or multiple encoding analysis in the 3BNC117 treated cohort (see Materials & Methods).

| hxb2_coord_AA | Participant | Method | Putative Escape |
|---|---|---|---|
| 90 | 2D1 | MPL | FALSE |
| 97 | 2E2 | multi_encodings | FALSE |
| 145 | 2C5 | multi_encodings | FALSE |
| 274 | 2E2 | MPL | TRUE |
| 279 | 2E3 | multi_encodings | TRUE |
| 279 | 2E7 | multi_encodings | TRUE |
| 282 | 2E4 | multi_encodings | TRUE |
| 282 | 2E4 | MPL | TRUE |
| 282 | 2E5 | multi_encodings | TRUE |
| 282 | 2E5 | MPL | TRUE |
| 295 | 2E2 | MPL | FALSE |
| 297 | 2E2 | MPL | FALSE |
| 364 | 2C5 | MPL | TRUE |
| 440 | 2E3 | MPL | FALSE |
| 459 | 2E7 | MPL | TRUE |
| 459a | 2C5 | MPL | TRUE |
| 459a | 2D1 | MPL | TRUE |
| 464 | 2E3 | MPL | TRUE |
| 706 | 2E1 | multi_encodings | FALSE |
| 754 | 2E7 | MPL | FALSE |

### 3BNC117 MPL additional variation sites

**Supplemental Table 10.**
Sites in 3BNC117 contact regions for which the independent locus version of MPL inferred selection coefficients $s \geq 0.044$, but the full version of MPL inferred lower selection coefficients when accounting for linkage to other sites. Columns 's_SL' and 's_MPL' hold the selection coefficients for the independent locus version of MPL and the full version of MPL, respectively.

| Participant | hxb2_coord_AA | s_MPL | s_SL |
|---|---|---|---|
| 2C5 | 279 | 0.041 | 0.0857 |
| 2C5 | 281 | 0.04 | 0.072 |
| 2D1 | 364a | 0.014 | 0.0669 |
| 2E1 | 460 | 0.027 | 0.0467 |
| 2E1 | 463a | 0.028 | 0.0485 |
| 2E2 | 461 | 0.002 | 0.0538 |
| 2E4 | 461 | 0.015 | 0.0678 |
| 2E7 | 279 | 0.035 | 0.0725 |
| 2E7 | 462 | 0.015 | 0.0626 |

#### 3BNC117 Newly discovered AAs at putative escape loci

**Supplemental Table 11.**
Amino acids which were found in SMRT-UMI sequences at loci under strong positive selection after 3BNC117 infusion but were not present in the original SGS sequencing data (Caskey et al., 2015; Schoofs et al., 2016) at time points after 3BNC117 infusion. The “Max Frequency” column indicates the maximum allele frequency the mutation reached and the “Max Timepoint” column indicates the time at which it reached this maximum. Note, mutation 459aG in participant 2D1 was detected by SGS at day 0, but not after 3BNC117 infusion.

| Participant | Escape Site | Unique AA | D0 | W1 | W12 | W4 | W8 | Max Timepoint | Max Frequency |
|---|---|---|---|---|---|---|---|---|---|
| 2C5 | 364 | T | | | | | 2 | W8 | 0.007462686567 |
| 2C5 | 459a | N | 2 | | 2 | 3 | 8 | W12 | 0.05405405405 |
| 2D1 | 459a | G | 12 | 17 | 47 | 60 | 33 | D0 | 1 |
| 2D1 | 459a | K | | 1 | 4 | 1 | 1 | W12 | 0.0701754386 |
| 2D1 | 459a | R | | | 1 | 3 | 1 | W4 | 0.03296703297 |
| 2E1 | 706 | K | | 1 | | 1 | 4 | W8 | 0.0350877193 |
| 2E2 | 274 | F | | | 4 | 1 | 3 | W8 | 0.25 |
| 2E2 | 295 | T | 1 | 1 | | | | W1 | 0.09090909091 |
| 2E3 | 279 | A | | | | 3 | 2 | W4 | 0.01181102362 |
| 2E4 | 282 | Q | | | 2 | 3 | 1 | W4 | 0.01111111111 |
| 2E4 | 282 | R | | | 4 | 3 | 1 | W12 | 0.01428571429 |

#### Sample ID Sequences

**Supplemental Table 12.**

Chosen to have maximal inter-index genetic distance

Sample Index sequences (16 chosen to have maximal inter-index genetic distances):

| | | | | | |
|---|---|---|---|---|---|
| 01 | ACAGTG | 07 | ACTGAT | 14 | GTCATC |
| 02 | CACTCA | 08 | TGACCA | 15 | CGAGTA |
| 03 | GGTAGC | 09 | GCTCAT | 16 | GACAGA |
| 04 | TAGCTT | 11 | CGATGT | 17 | TAGAGC |
| 05 | CTATAC | 12 | ATGCTG | | |
| 06 | ATCACG | 13 | ACGATC | | |

Primer Sequences used for SMRT-UMI protocol (Westfall et al., 2024)

**Supplemental Table 13.**

| Primer Name | Use | Sequence |
|---|---|---|
| PBX_R9165 | SMRT-UMI Subtype B cDNA primer | CCCGCGTGGCCTCCTGAATTATCCGCTCCGTCCGACGACTCACTATAXXXXXXNNNNNNNNCTGGTGTGTARTTYTGCCAATCAG |
| PB_AX_nef67_degen | SMRT-UMI Alternate cDNA primer | CCCGCGTGGCCTCCTGAATTATCCGCTCCGTCCGACGACTCACTATAXXXXXXNNNNNNNNGGTCTTAAAGGYACCTGAGGTCTGACTGGAAAGCC |
| PB_AX_nef67_degen_mod | SMRT-UMI Alternate cDNA primer | CCCGCGTGGCCTCCTGAATTATCCGCTCCGTCCGACGACTCACTATAXXXXXXNNNNNNNNGGTCTTAAAGGYACCTGAGGTGTGACTGGAAAACC |
| R9165 | SGA Subtype B cDNA and PCR 1 Reverse primer | CTGGTGTGTARTTYTGCCAATCAG |
| F5876 | SMTR-UMI and SGA PCR 1 Forward primer | TAGAGCCCTGGAAGCATCCAGGAAG |
| F5982A | SMTR-UMI PCR 2 Forward primer | TAGGCATCTCCTATGGCAGGAAGAAG |
| F5066alt1 | SMTR-UMI Alternate PCR 1 Forward primer | TATGGAAAACAGATGGCAGGTGMTGRT |
| F5088alt1 | SMTR-UMI Alternate PCR 2 Forward primer | GATTGTGTGGCARGTAGACAGRATG |
| PB-R1-alt1 | SMTR-UMI PCR 1 Reverse primer | CCCGCGTGGCCTCCTGAATTAT |
| PB-R2-alt1 | SMTR-UMI PCR 2 Reverse primer | CCGCTCCGTCCGACGACTCACTATA |

| | | |
|---|---|---|
| envB5in-tag | SGA PCR 2 Forward Primer | TCGTCGGCAGCGTCTTAGGCATCTCCTATGGCAGGAAGAAG |
| envB3in-tag | SGA PCR 2 Reverse Primer | GTCTCGTGGGCTCGGGTCTCGAGATACTGCTCCCACCC |
| *XXXXXX = 6 base pair sample ID (supplemental table 11) | | |
| *NNNNNNNN = 8 base pair random nucleotides | | |
