## Supplemental Methods for "Recurrent mutations drive rapid HIV escape from two broadly neutralizing antibodies *in vivo*"

### Extended Description of RNA isolation and cDNA synthesis

Frozen plasma samples (1mL) were thawed and viral RNA isolated using MinElute Virus Spin kits (Qiagen, #55704) or Viral RNA Mini spin kits (Qiagen, #52904). cDNA synthesis performed from viral RNA using HIV-1 specific oligos or SMRT-UMI cDNA 5' ultramers comprised an HIV-1 specific binding region, an 8bp random nucleotide UMI, a 6bp sample ID (Supp. Table 12), and two reverse primer binding regions (Westfall et al., 2024). cDNA was synthesized using SuperScript IV Reverse Transcriptase (10U/uL, Thermo Fisher Scientific), supplemented with SUPERaseIn RNAse Inhibitor (1U/uL, Thermo Fisher Scientific), ThermaStop RT (0.2U/uL, Millapore Sigma, discontinued) and standard or SMRT-UMI oligos (0.4uM, IDT). cDNA reactions were incubated for 1 hour at 50C, then inactivated at 80C for 10 minutes. RNA was removed with RNase H for 20 min at 37C (0.1U/uL, Thermo Fisher Scientific) and cDNA purified using RNAClean XP beads (Beckman Coulter) at a 1:1 ratio and washed 3x with 80% ethanol.

### Extended Description of Amplification

HIV env cDNA concentration was estimated using an end point dilution 25uL nested PCR using PrimeSTAR GXL polymerase (1.25U/uL, Takara Bio Inc), a 1uL cDNA input, and 0.25uM forward and reverse PCR 1 primers as previously described (Westfall et al., 2024). End point nested PCR was performed at the following cycling conditions for PCR 1: 2min at 98C; 35 cycles of 10 sec at 98C, 15 sec at 60C, and 3:15min at 68C; 7 min at 68C; 4C hold. The following cycling conditions were performed for PCR 2 with a 1uL PCR 1 input and forward and reverse PCR 2 primers: 2min at 98C; 35 cycles of 10 sec at 98C, 15 sec at 62C, and 3:15min at 68C; 7 min at 68C; 4C hold. Positive end point dilution reactions were defined as a 3kb band visualized by gel electrophoresis (Rodrigo et al., 1997) and used to estimate cDNA concentration using the Quality webtool (https://quality.fredhutch.org/).

Samples with low viral load, where cDNA was synthesized without SMRT-UMI oligos, were amplified at limiting dilution, to ensure positive PCRs contained only a single cDNA template, according to the PCR conditions listed above. PCR 2 primers contained 5' tag sequences complementary to unique index primers added during a third PCR to allow for pooling of positive PCRs for sequencing.

For SMRT-UMI sequencing, an estimated 50 HIV env cDNA molecules were input into each 25uL PCR reaction and were amplified using 0.25uM forward and reverse PCR 1 primers at the following cycling conditions: 2min at 98C; 20 cycles of 10 sec at 98C, 15 sec at 60C, and 3:15min at 68C; 7 min at 68C; 4C hold. First-round PCR product was pooled and cleaned by magnetic bead size selection (AMPure XP, 0.5x) and a second-round PCR was performed using 1uL of bead cleaned 1st round product and 0.25uM second round PCR primers at the following cycling conditions: 2min at 98C; 20 cycles of 10 sec at 98C, 15 sec at 62C, and 3:15min at 68C; 7 min at 68C; 4C hold.
A heteroduplex resolution step was conducted at the end of the second PCR, which consisted of spiking in PrimeSTAR GXL, 2nd round forward and reverse primers and dNTPs to each reaction tube and incubating for: 2min at 98C; 15s at 62C; 10min at 68C; and a 4C hold. Final

amplified products were confirmed by a 3kb band visualized by gel electrophoresis and purified by magnetic bead size selection (AMPure XP magnetic beads, 0.7x).

All cDNA and PCR primers ordered from IDT and listed in Supp. Table 13. Index primers listed along with demultiplexing pipeline at https://github.com/lcohnlab/sga-pipeline.
